# Chemical Probe Discovery for DEAD-box RNA Binding Protein DDX21 using Small Molecule Microarrays

**DOI:** 10.1101/2025.04.18.649545

**Authors:** Toshihiko Aiba, Eliezer Calo, Angela N. Koehler

## Abstract

The DEAD-box family of ATPases plays a critical role in nearly all stages of RNA metabolism, from transcription to degradation, and serves as a major regulator of biomolecular condensates. Dysregulation of DEAD-box proteins is well established in a variety of diseases, including cancer and neurodegenerative disorders, making them attractive therapeutic targets. However, their classification as “undruggable” has historically hindered small molecule-based modulation. In this study, we focus on DDX21, a member of the DEAD-box family involved in ribosome biogenesis and transcriptional regulation. As a proof of concept for targeting such RNA-binding proteins, we developed a lysate-based small molecule microarray platform to identify compounds that directly bind DDX21. This screen led to the discovery of KI-DX-014, a small molecule compound capable of inhibiting DDX21 interaction with RNA. KI-DX-014 modulated RNA-dependent functions of DDX21, including its ATPase activity and biomolecular condensate formation. Furthermore, KI-DX-014 attenuated the DDX21-dependent release of P-TEFb from the 7SK snRNP complex in vitro, suppressed P-TEFb-dependent phosphorylation of the RNA polymerase II CTD, and induced developmental defects in zebrafish embryos. These findings reveal a previously unexploited therapeutic avenue and establish KI-DX-014 as a valuable chemical probe for dissecting the biological functions of DDX21 in both normal physiology and disease states.

## Introduction

DDX21 is a member of the DEAD-box protein family, capable of binding and remodeling RNA structures and RNA-protein complexes through ATP hydrolysis^1,2^. DDX21 is involve in multiple biological processes, including ribosome biogenesis^3,4^, RNA polymerase II (Pol II) elongation^3^, and tissue differentiation^5,6^. DDX21 achieves these functions by interacting with process-specific RNA substrates such as the pre-ribosomal RNA, snoRNAs, and 7SK snRNA^3^. DDX21 RNA substrates stimulate its ATPase activity^2,7,8^.

During RNA Pol II transcription, DDX21 facilitates the release of positive transcription elongation factor b (P-TEFb) from its inhibitory association with the 7SK ribonucleoprotein complex (7SK snRNP)^3,9^. Once released, P-TEFb is activated and phosphorylates multiple substrates, including the C-terminal domain (CTD) of Pol II, thereby promoting transcription elongation^10^. Additionally, DDX21 can undergo liquid-liquid phase separation (LLPS) *in vitro* and in cells via its RNA-binding domain and intrinsically disordered regions^11,12^. These LLPS activities of DDX21 has been associated with DDX21 cellular localization^12^, transcription control^13,14^, and regulation of gene expression^15^. Collectively, these diverse functions of DDX21 underscore the critical roles of this DEAD-box protein in development^16^, congenital diseases^17,18^ and various cancers^15,18–21^. Despite increasing evidence supporting the involvement of DDX21 in various cellular processes and diseases, the development of chemical probes targeting this protein has lagged, limiting our ability to elucidate the connection between its diverse functions and disease mechanisms. Therefore, the development of novel, specific chemical probes targeting DDX21 is essential to drive further advancements in this field.

Chemical probes are valuable tools that can provide insights into the biological functions of target proteins through the pharmacological modulation of their targets^22^. However, there are no chemical probes that specifically target DDX21. RNA-binding proteins (RBPs), such as DDX21, are considered undruggable targets^23,24^. This is primarily due to DDX21’s highly conserved ATPase domains, shared among all DEAD-box proteins, its lack of distinct small-molecule binding pockets, and the limited feasibility of functional high-throughput assays.^24^. Like most RBPs, DDX21 contains conformationally dynamic regions, often referred to as intrinsically disordered regions (IDRs), that mediate diverse molecular functions such as protein-protein interactions, forming ribonucleoprotein complexes, or driving LLPS within cells^25–28^. Reconstituting these IDR-driven activities in vitro with purified proteins presents significant challenges, which, in turn, limits the feasibility and accuracy of biochemical assays. Nonetheless, recent advances in small-molecule screening techniques and an understanding of the biology of RBPs have enabled the identification of functional chemical probes for selected RBPs^29,30^.

Given the challenges of targeting RBPs, such as DDX21, we utilized small molecule microarrays (SMMs) to discover small-molecule binders of DDX21. This method enables screens for binders to proteins residing in cell lysates, likely reflecting a protein state that is more like a native state relative to purified protein^31^, as it preserves the network of protein–protein and protein–RNA interactions. This screening platform uses small molecule compounds printed on a glass surface for the fluorescent detection of protein-small molecule interactions^32,33^. To date, this method has been employed to discover small molecules that bind to diverse targets, including the transcription factor ETV1^34^, the MAX homodimer^35^, and CDK9 derived from and AR-V interactome-focused screen^36^. Here we introduce a lysate-based SMM screening approach that facilitated the discovery and characterization of KI-DX-014, a tool compound that inhibits DDX21’s interaction with RNA by targeting its intrinsically disordered C-terminal domain. KI-DX-014 effectively disrupts RNA-dependent transcriptional activities of DDX21 both *in vitro* and *in vivo*. This approach provides a powerful and versatile platform for the discovery of small molecule modulators targeting other RBPs.

## Results

### SMM screening identifies small molecule binders of DDX21

We used small molecule microarrays (SMMs) to screen nuclear extracts from HEK293 cells expressing a fluorescence-labeled N-terminal HaloTag®^37,38^ fused to full-length DDX21 (Halo-DDX21) against a library of 115,805 commercially available small molecules (Fig. 1a, b). In this assay, the target protein, Halo-DDX21, was labeled in situ with a Halo-Tag fluorescent ligand, which minimized the washing steps. First, HEK293 cells stably expressing tetracycline-inducible Halo-DDX21 or Halo-GFP (control) were generated using the PiggyBac transposon vector system. Transgenes were expressed by the addition of doxycycline (DOX) for 48 h, and nuclear extracts were prepared for SMM screening. The SMM microarray was first incubated with the nuclear extract containing fluorescence-labeled Halo-DDX21 for target screening or Halo-GFP for control screening. Following this, the SMM microarray was washed, imaged, and fluorescent intensity for protein binding to each small molecule feature was measured. The signal-to-noise ratio (SNR) of each printed spot was calculated by dividing the foreground signal by the background signal, and then normalizing the SNR by calculating a Robust Z-Score^39^. Compounds were considered assay positives if their Robust Z-score was greater than 3 across all four replicates as previously reported^38^. To remove false positives, the compounds were manually inspected for spot fluorescence and positives in the Halo-GFP control screening were eliminated. In total, 583 compounds were identified as putative DDX21 and interactome binders with a hit rate of 0.5% (Fig. 1a, c).

**Figure 1:**
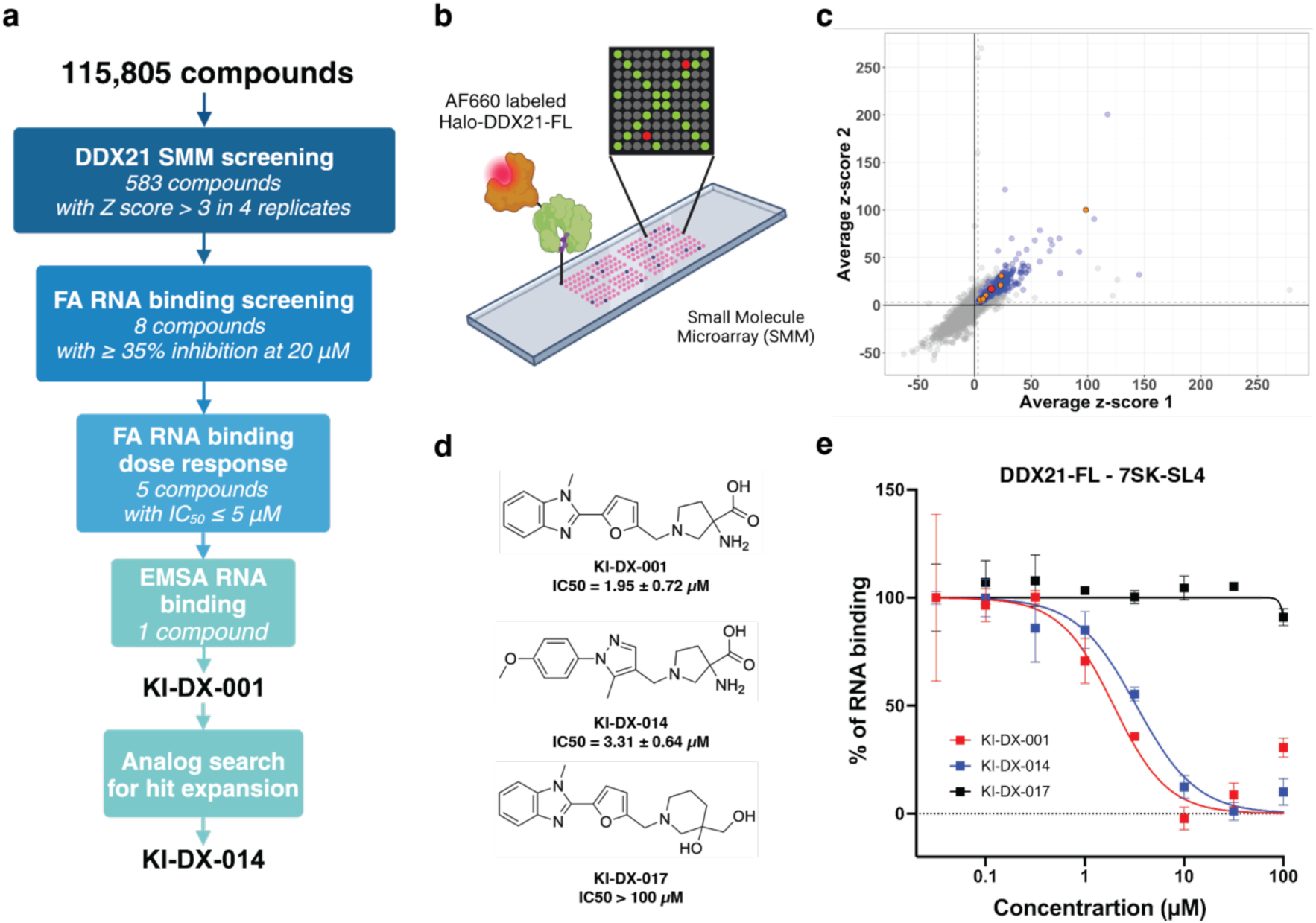
Discovery of a DDX21 small molecule binder, KI-DX-014 as a DDX21-RNA interaction inhibitor. **a**. Prioritizing scheme for DDX21 binders that modulate protein-RNA interaction leading to hit probe candidate KI-DX-001 and KI-DX-014 as an advanced probe candidate. SMM, small molecule microarray; FA, fluorescence anisotropy; EMSA, electrophoretic mobility shift assay. **b**. Schematic of SMM assay design. SMM positives were detected by Alexa Fluor 660-labeled N-terminal HaloTag® fused DDX21 in lysates. Created with BioRender.com. **c**. Screening result showing binding of 115,805 compounds to Halo-DDX21 on SMM. Scatterplot showing the average robust Z-scores of the signal/background ratio for each microarray slide. SMM hit compounds with robust Z-score > 3 in 4 replicates and signal/background ratio > 1.3 in 4 replicate and excluded hit compounds from control SMM screening using Halo-GFP are highlighted in blue. Validated compounds for which dose-response experiments in the RNA binding inhibition assay were performed are highlighted in orange; KI-DX-001 is highlighted in red. **d**. Chemical structures of KI-DX-001, KI-DX-014 and negative probe, KI-DX-017. **e**. Fluorescence anisotropy assay for competition of 5’-FAM labeled 7SK-SL4 RNA for binding to DDX21-FL using KI-DX-001 (red), KI-DX-014 (blue) and KI-DX-017 (black). Data are shown as mean values ± s.d. for three technical replicates.

### Secondary screening of DDX21 binders that inhibit RNA binding via fluorescence anisotropy

DDX21 is a multifunctional protein that engages in diverse cofactor-dependent activities, predominantly facilitated by its RNA-binding properties^40^. To identify DDX21 RNA-binding inhibitors from SMM screening hits, we developed a fluorescence anisotropy assay to systematically and quantitatively assess DDX21’s RNA-binding activity. First, purified full-length DDX21 was prepared from HEK293 cell lysates expressing Halo-DDX21. Halo-DDX21 was immobilized on magnetic beads coated with HaloTag® ligand. The DDX21 protein was released from the magnetic beads by cleaving a TEV linker, positioned between the Halo-tag and DDX21 coding sequence, using HaloTEV protease. The protease remained bound to the magnetic beads via the HaloTag®, resulting in a highly pure DDX21 protein sample (Supplementary Fig. 1a). Subsequently, we utilized various fluorescently labeled RNA substrates of DDX21 (Supplementary Figure 1b) to develop a fluorescence anisotropy-based RNA binding assay. This assay was used to characterize the biochemical and functional properties of the eluted DDX21 protein as well as the initial SMM-positive compounds. Among the RNA substrates tested, two stem loops of 7SK snRNA, 7SK-SL1 and 7SK-SL4, both endogenous targets of DDX21, demonstrated a pronounced anisotropy shift even at low nanomolar concentrations (Supplementary Figure 1b). In contrast, a recently identified RNA oligonucleotide substrate of DDX21 exhibited only a modest change in anisotropy values under similar conditions^5^ (Supplementary Figure 1b). We then determined the binding affinity of 7SK-SL1 and 7SK-SL4 to DDX21 (Supplementary Fig. 1c, d). Both RNA probes demonstrated strong binding to DDX21 and were used for subsequent secondary screening.

To evaluate RNA-binding inhibition, we conducted secondary screens using the SMM assay positives at a fixed concentration of 20 µM. Compounds were classified as active if they reduced RNA binding to less than 50% relative to the DMSO control (Supplementary Figure 2a). To assess the reproducibility of the selected active compounds, triplicate assays were conducted. We identified eight compounds that consistently inhibited DDX21’s ability to bind RNA, reducing RNA binding to less than 35% (Supplementary Figure 2b). We purchased seven of the eight reproducibly active compounds and systematically evaluated their dose-dependent RNA-binding inhibition (Supplementary Figure 2c). Notably, some compounds (i.e., compounds 17, 18 and 20) lost their RNA binding inhibitory activity at higher concentration ranges due to their poor solubility (Supplementary Fig. 2c). Ultimately, four compounds consistently exhibited RNA-binding inhibition with IC50 values below 5 µM (Supplementary Figures 1d and 2d).

To further validate the RNA-binding inhibition of the selected compounds, we developed an orthogonal infrared dye-based electrophoretic mobility shift assay (IR-EMSA). First, we confirmed that the binding affinity of the IRDye 800CW-conjugated 7SK-SL4 to DDX21 was comparable to the affinity observed in the fluorescence anisotropy assays (Supplementary Figure 1d). Next, we assessed selected hits for RNA-binding inhibition by IR-EMSA. Among these, KI-DX-001 demonstrated notable RNA-binding inhibition, reducing binding by 35% at a concentration of 20 µM (Supplementary Figure 2d).

### Hit expansion by Structure-Activity Relationships

To assess structure-activity relationships (SARs) around KI-DX-001, we performed a 2D similarity search using the commercially available compound database ZINC20^41^. Based on their scoring, chemical structure, and availability, we selected eleven compounds (Supplementary Fig. 3a). We then evaluated the ability of these compounds to inhibit RNA binding using fluorescence anisotropy (Supplementary Fig. 3b). We first explored the impact of substituting the cyclic amino acid moiety of KI-DX-001. Although analogues containing triazole (KI-DX-009), sulfone amide (KI-DX-010) and morpholine amide (KI-DX-013) exhibited the RNA-binding inhibitory activity (Supplementary Fig. 3a), the significant reductions in RNA-binding inhibitory activity (IC50 > 100 µM) were observed for analogues bearing morpholine-2-carboxylic acid amide (KI-DX-007), imidazole (KI-DX-008) (Supplementary Fig. 3a) and diol (KI-DX-017) (Fig. 1d, e). Subsequently, we examined the substitution of the core heterocyclic structure of KI-DX-001. While the thiazole analogue (KI-DX-011) was inactive in the RNA binding inhibition, analogues including oxadiazole, pyrazole, phenyl and imidazole (KI-DX-012, 014, 015 and 016) retained inhibitory activity. Among the evaluated compounds, KI-DX-014 showed high RNA-binding inhibitory activity (IC50 3.31 ± 0.64 µM) (Fig. 1d, e) and exhibited dose-dependent RNA-binding inhibition in an EMSA assay (Supplementary Fig. 3c). From our initial SAR experiment relying on commercial compounds, we selected KI-DX-014 for further validation because the compound retained RNA binding inhibitory activity at high doses (Fig. 1e), has better solubility in aqueous solution, and showed more pronounced RNA binding inhibition in an EMSA assay (Supplementary Fig. 3c) compared to KI-DX-001.

### KI-DX-014 directly binds to DDX21 and inhibits DDX21-RNA interaction via the C-terminal intrinsically disordered domain

To obtain evidence for the direct interaction between DDX21 and KI-DX-014, we purified a histidine-tagged DDX21 construct containing the RNA-binding domains, specifically the helicase core and C-terminal domain, referred to as DDX21-HC+CTD. We then assessed its binding affinity for 7SK-SL4 RNA and found it to be comparable to the full-length protein (Fig. 2a, Supplementary Fig. 4a, b). We used microscale thermophoresis (MST)^42^ to analyze the affinity of KI-DX-014 for DDX21-HC+CTD (*K_d_* = 15.9 ± 3.9 µM) (Fig. 2b). This result suggests that KI-DX-014 inhibits RNA binding through direct binding to DDX21.

**Figure 2:**
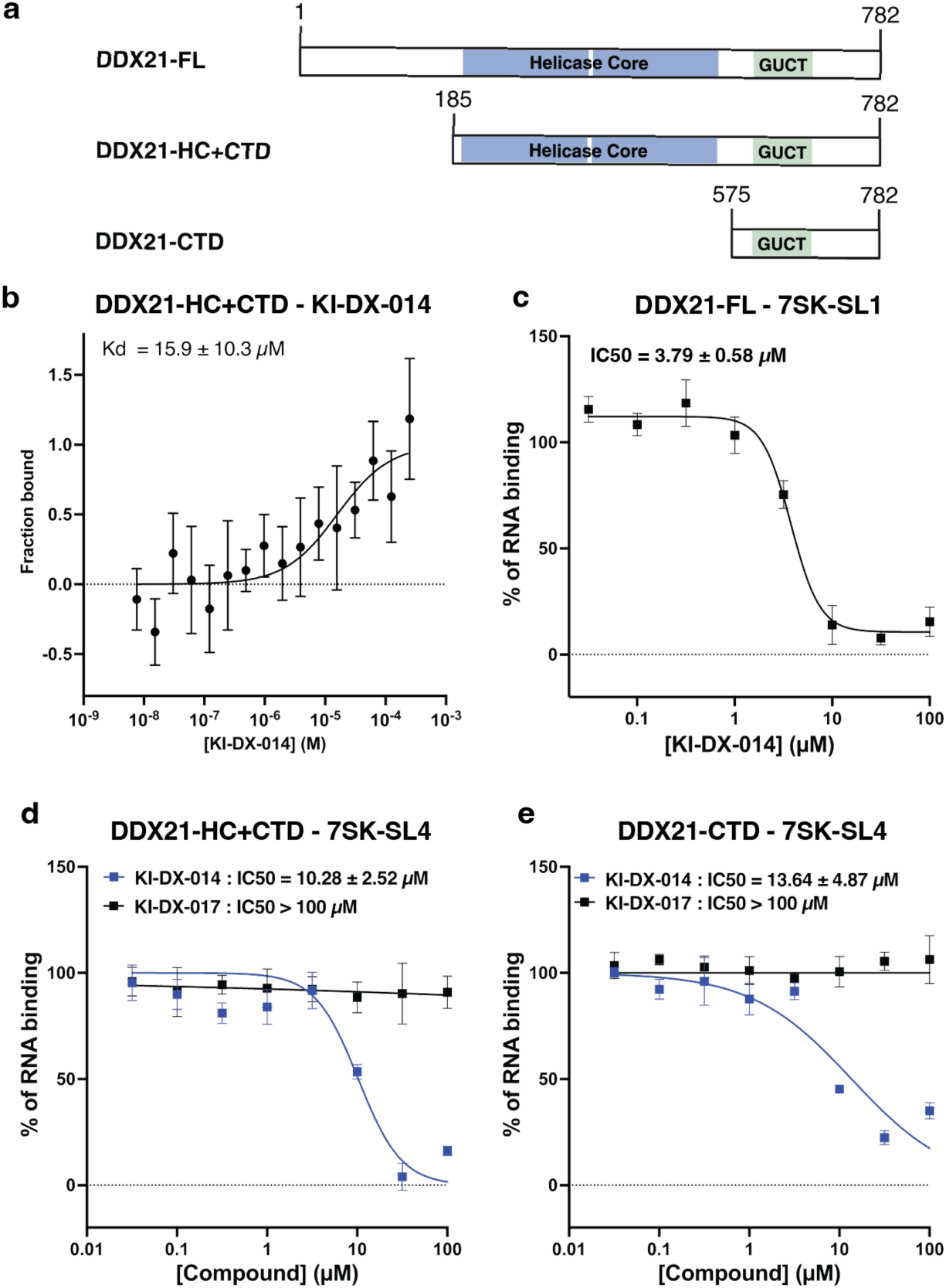
KI-DX-014 directly binds to DDX21 and inhibits DDX21-RNA interaction via the C-terminal domain (CTD). **a**. Schematic representation of DDX21-FL, the full length DDX21 and truncation constructs of DDX21, DDX21-HC+CTD and DDX21-CTD used in this study. Created with BioRender.com. **b**. Microscale thermophoresis (MST) interaction analysis of Red-tris-NTA 2nd generation dye labeled N-terminal His-Tag DDX21-HC+CTD against KI-DX-014. Data are shown as mean values ± s.d. for four biological replicates. **c**. Fluorescence anisotropy assay for competition of 5’-FAM labeled 7SK-SL1 RNA for binding to DDX21-FL using KI-DX-014. Data are shown as mean values ± s.d. for three technical replicates. **d**. Fluorescence anisotropy assay for competition of 5’-FAM labeled 7SK-SL4 RNA for binding to DDX21-HC+CTD using KI-DX-014. Data are shown as mean values ± s.d. for three technical replicates. **e**. Fluorescence anisotropy assay for competition of 5’-FAM labeled 7SK-SL4 RNA for binding to DDX21-CTD using KI-DX-014. Data are shown as mean values ± s.d. for three technical replicates.

Next, we characterized the interaction between DDX21 and RNA, as well as the inhibitory activity of KI-DX-014. We found that KI-DX-014 also inhibits the binding of DDX21 to 7SK-SL1, another stem-loop region of 7SK, which DDX21 binds with similar kinetics to 7SK-SL4 (Fig. 2c). Previous study has shown that the C terminal domain of DDX21 is RNA binding domain, facilitating the binding of DDX21 to highly structured RNAs, such as the ribosomal RNA^12^ and G4 quadruplexes^43^. We next purified DDX21-CTD (Fig. 2a) and analyzed its affinity for 7SK-SL4 using fluorescence anisotropy and EMSA (Supplementary Fig. 4c–e). The results showed that 7SK-SL4 binds to DDX21-CTD with an affinity comparable to that of the full-length protein (Supplementary Fig. 4d), confirming that the CTD is a domain of DDX21 capable of binding to structured RNAs.

To examine whether KI-DX-014 inhibits RNA binding by engaging with DDX21’s RNA-binding domain, we performed competition experiments between KI-DX-014 and 7SK-SL4 using both DDX21-HC+CTD and DDX21-CTD constructs. KI-DX-014 effectively competed with 7SK-SL4, suggesting that both bind to the same site (Fig. 2d, e). To test specificity, we performed the experiment with a structurally similar compound, KI-DX-017, which lacked RNA-binding inhibitory activity against full-length DDX21. As expected, KI-DX-017 was unable to compete with 7SK-SL4 (Fig. 2d, e).

Next, we evaluated the binding of a non-structured RNA, U15 RNA, to DDX21-FL. Non-structured RNA can interact non-specifically with the helicase core domain of DDX21^7,44^. While U15 showed binding to DDX21-FL (Supplementary Fig. 4f), its affinity was approximately 50-fold lower than that of the endogenous structured RNAs 7SK-SL1 and 7SK-SL4 (Supplementary Fig. 1c). We then investigated the inhibitory effect of KI-DX-014 on U15 binding to DDX21-FL. KI-DX-014 exhibited no inhibitory activity against U15 binding (Supplementary Fig. 4g), likely because U15 preferentially interacts with the helicase core domain of DDX21. Consistent with this, KI-DX-014 does not impair the ability of the DDX21 to unwind an RNA duplex (Supplementary Figures 4h, i), suggesting that KI-DX-014 specifically targets and affects the RNA-binding activities of the DDX21 C-terminal domain. Taken together, these results indicate that KI-DX-014 specifically inhibits the binding of structured RNAs, such as 7SK-SL1 and 7SK-SL4, which interact with the C-terminal domain of DDX21. In contrast, it does not inhibit the unwinding activity of DDX21 or its binding to non-structured RNAs, which likely interact with the helicase domain.

### KI-DX-014 modulates RNA binding-dependent ATPase activity of DDX21

The ATPase activity of DDX21 depends on its ability to bind RNA, which occurs through specific conformational states^7,44^. To test whether KI-DX-014-mediated inhibition of RNA-binding impacts DDX21 ATPase activity, we measured ATP hydrolysis with the ADP-Glo assay^45^. The presence of 7SK-SL4 significantly enhanced the ATP hydrolysis activity of DDX21 (Fig. 3a). This activation was effectively inhibited by KI-DX-014 in a dose-dependent manner (Fig. 3a), with a reduction of maximal ATP hydrolysis activity (Fig. 3a). We have shown that KI-DX-014 does not alter unwinding activity of double strand RNA which is driven by ATP hydrolysis (Supplementary Figures 4h, i). Taken together, these findings suggest that KI-DX-014 does not inhibit ATP, the substrate of ATPase, and thus noncompetitively attenuates the ATPase activity of DDX21. As expected, U15 also stimulated DDX21 ATPase activities (Fig. 3b). However, the magnitude of ATPase activation elicited by U15 was less pronounced than that observed with 7SK-SL4 binding (Fig. 3a, b), consistent with U15 low binding affinity to DDX21 (Supplementary Fig. 4f) when compared to 7SK-SL4 (Supplementary Fig. 1c). Surprisingly, KI-DX-014 also exhibited dose-dependent inhibition of DDX21 ATPase activation induced by U15 (Fig. 3b). These results suggest that KI-DX-014 binding to DDX21 not only inhibits the interaction with structured RNAs but also prevents the allosteric regulation of DDX21 ATPase activity mediated by its C-terminal domain. Thus, KI-DX-014 allosterically modulate DDX21 ATPase activity.

**Figure 3:**
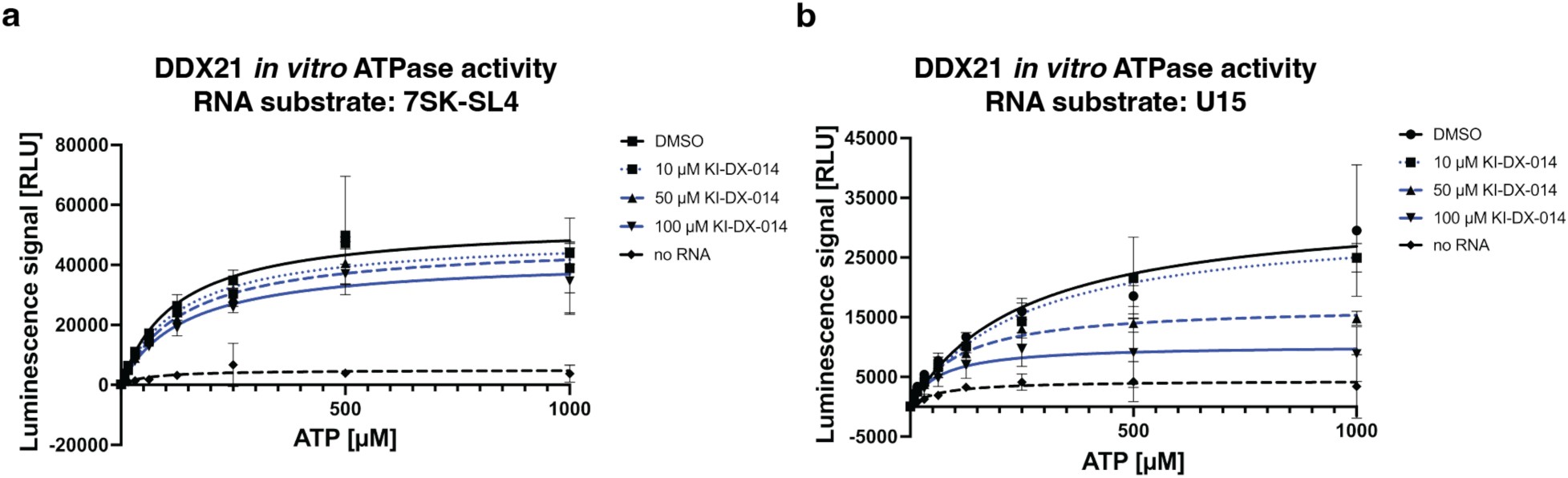
KI-DX-014 regulates RNA binding-dependent ATPase activity **a**,**b**. Effects of KI-DX-014 on DDX21 ATPase activity using 7SK-SL4 (**a**) or U15 (**b**) as an RNA substrate in the presence of various concentrations of ATP. ATP was used at concentrations ranging from 7.81 to 1000 µM and ATPase reaction was performed for 1 h. Data are shown as mean values ± s.d. for three technical replicates.

### KI-DX-014 regulates RNA binding-dependent liquid-liquid phase separation of DDX21

DEAD-box RNA binding protein, including DDX21, undergo LLPS *in vitro* and modulate biomolecular condensates formation and activity in cells through their intrinsically disordered regions and interactions with other co-factors, including RNA^46^. A previous study demonstrated that RNA nucleates protein-RNA condensates and act as an architectural element that influences the composition and the morphology of biomolecular condensates^25^. Interestingly, the cellular localization of DDX21 is influenced, at least in part, by the ability of its C-terminal domain to undergo LLPS^12^. To examine whether KI-DX-014 disrupts DDX21-RNA interactions within biomolecular condensates and regulates the phase separation state driven by these interactions in vitro, we utilized a droplet formation assay with fluorescently labeled 7SK-SL4 and full-length DDX21 (Fig. 4a). We observed droplet formation when DDX21-FL is incubated with 10% PEG-8000 and fluorescently labeled 7SK-SL4 (Fig. 4b). Importantly, droplet formation is strictly dependent on the 7SK-SL4 RNA as treatment of the reaction with RNase abrogate droplet formation (Fig. 4b). The addition of KI-DX-014 led to a dose-dependent reduction in the size of these droplets (Fig. 4b, c). We observed a similar effect on droplets formed by U15 upon treatment with KI-DX-014 (Supplementary Fig. 5a, b). These data show that inhibition of DDX21-RNA binding by KI-DX-014 can modulate DDX21-RNA interactions within droplets and the ability of DDX21 to participate in LLPS *in vitro*.

**Figure 4:**
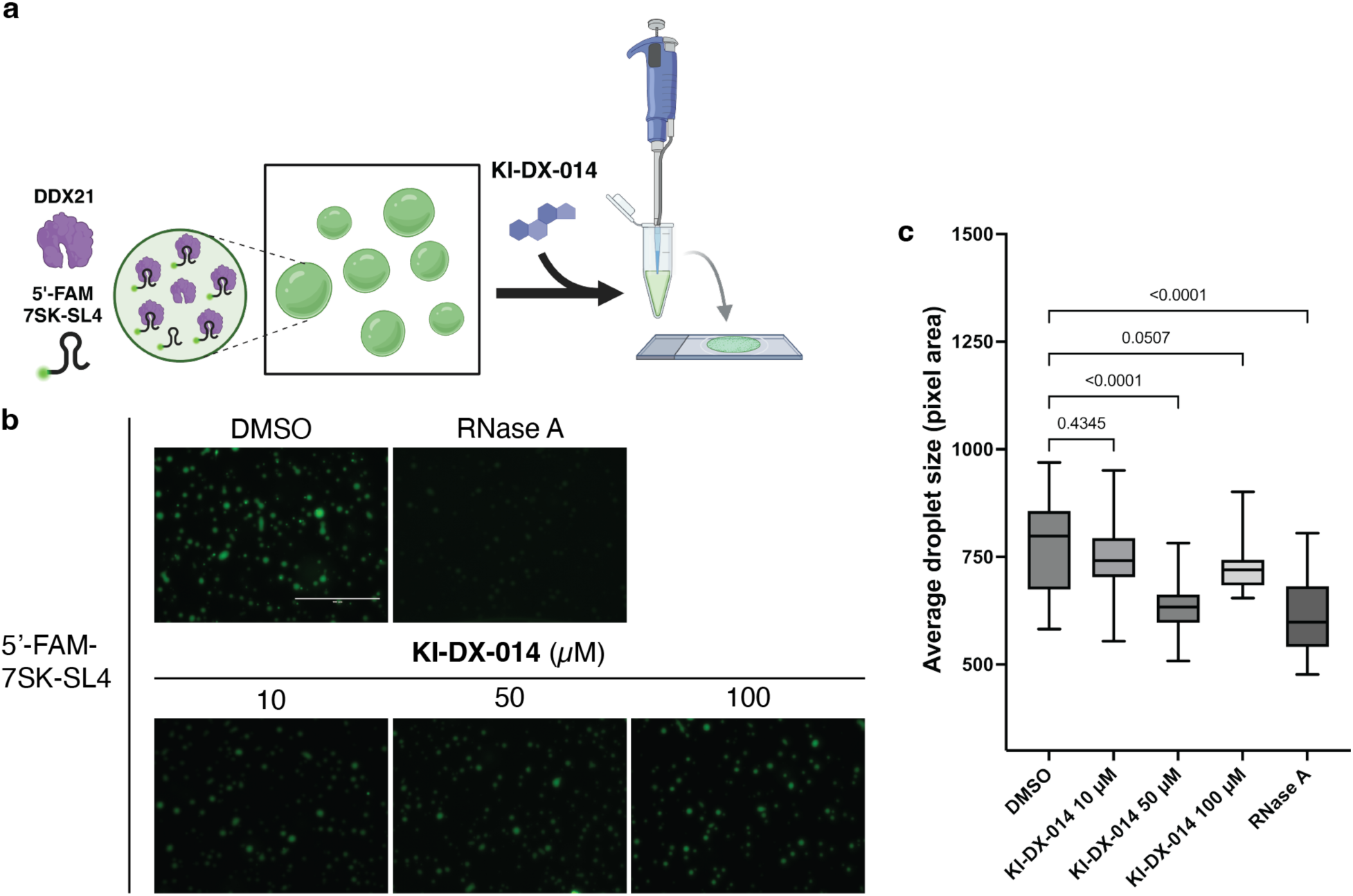
KI-DX-014 modulates protein-RNA phase separation *in vitro*. **a**. Schematic of *in vitro* DDX21 phase separation assay in the presence of KI-DX-014. Created with BioRender.com. **b**. Concentration dependent effects of KI-DX-014 on the morphological properties of DDX21 condensates, showing representative fluorescence images with 5’-FAM labeled 7SK-SL4 in the presence of KI-DX-014 or RNase A in 10% PEG8000; scale bar, 100 µm. **c**. Quantification of the average size of the droplets. Data are shown using box–whisker plots and error bars represents s.e.m.. *P* values were determined by one-way ANOVA with Dunnett’s test for multiple comparison.

### KI-DX-014 decreases release of P-TEFb from the 7SK snRNP

We then examined whether KI-DX-014 impacts more complex RNA-dependent biochemical processes governed by DDX21, such as the release of P-TEFb from the native 7SK ribonucleoprotein complex. P-TEFb is a heterodimer composed of CDK9 and Cyclin T1^47,48^. A major function of P-TEFb is to facilitates transcriptional elongation by phosphorylating Pol II pausing-inducing factors (DSIF and NELF) and the Pol II CTD— primarily, but not exclusively—at serine 2 (CTD S2P)^49^. P-TEFb is inactive when in complex with HEXIM1 and the 7SK snRNA and is released from this complex by release factors, most notably BRD4^50,51^. DDX21 has been shown to interact with the 7SK snRNP^3,52^ and to facilitates the release of P-TEFb in an ATPase activity-dependent manner^3,53^. To determine whether KI-DX-014 can regulate P-TEFb release by DDX21, we employed a P-TEFb release assay by purifying the 7SK snRNP via HEXIM1 immunoprecipitation^3,54^ (Fig. 5a). In the presence of ATP, purified DDX21 effectively released P-TEFb from the 7SK (Fig. 5b). Incubation with a non-hydrolysable ATP analog (AMPPCP) attenuated P-TEFb release, confirming that this activity requires DDX21 ATPase activity (Fig. 5b). Treatment with KI-DX-014 resulted in a dose-dependent decrease in P-TEFb release (Fig. 5b). This result indicates that the association of DDX21 with 7SK snRNP via RNA binding is necessary for P-TEFb release.

**Figure 5:**
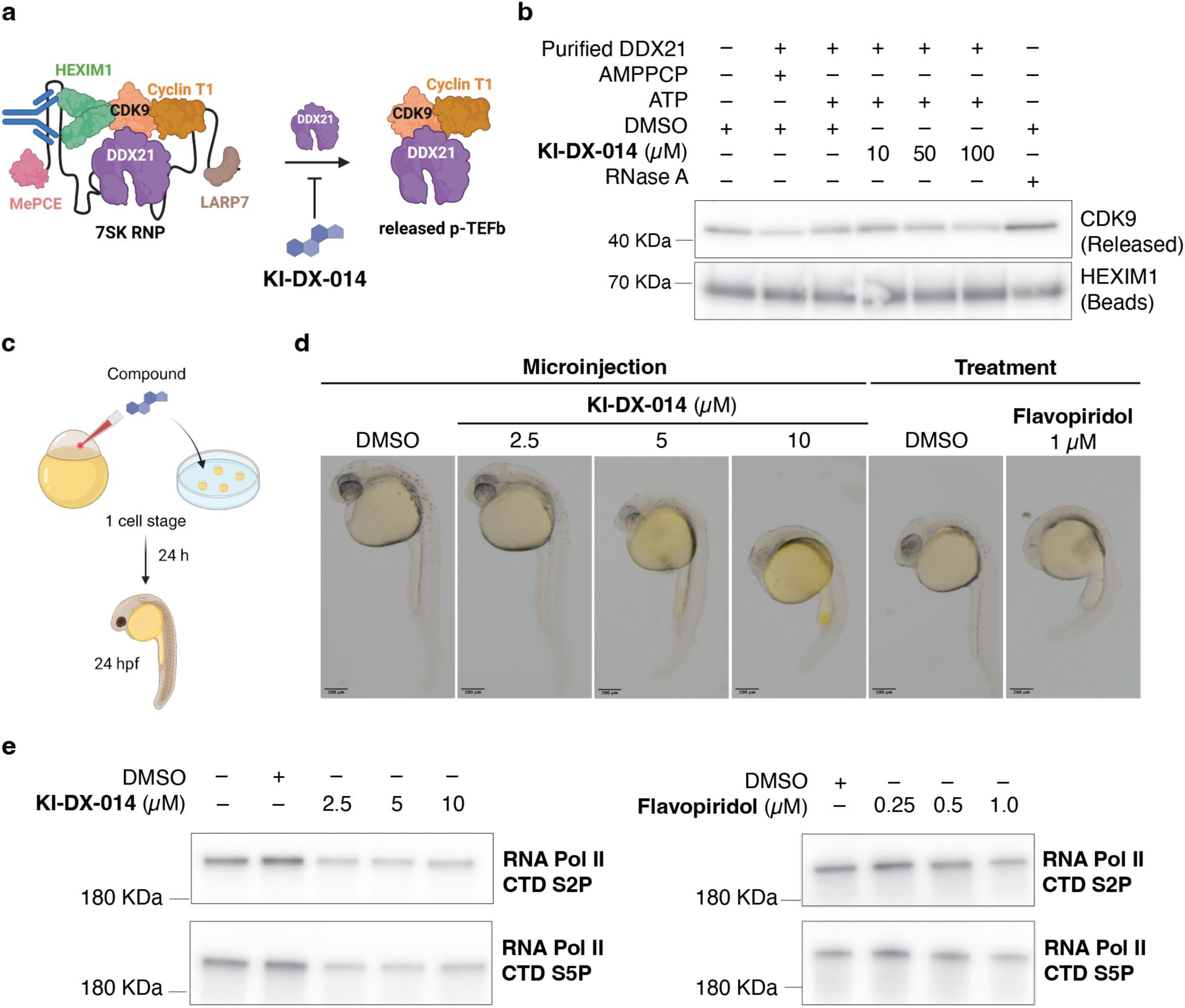
KI-DX-014 blocks release of P-TEFb from 7SK snRNP and causes developmental defects in zebrafish via Pol II phosphorylation. **a**. Schematic of P-TEFb release assay in the presence of KI-DX-014. Created with BioRender.com. **b**. Western blot analysis of released P-TEFb from HEXIM1 immunopurified 7SK snRNP upon addition of purified DDX21-FL and ATP in the presence of KI-DX-014. **c**. Schematic of *in vivo* treatment in zebrafish embryo model. KI-DX-014 or vehicle (DMSO) were microinjected to fertilized zebrafish embryos at 1 cell stage. Reference CDK9 inhibitor (Flavopiridol) or vehicle (DMSO) containing medium were treated to fertilized zebrafish embryos at 1 cell stage. After 24 h, zebrafish embryos were dechorionated for imaging and total protein extraction. Created with BioRender.com. **d**, Representative images of zebrafish embryos at 24 h.p.f. treated with KI-DX-014 and reference CDK9 inhibitors; scale bar, 200 µm. **e**, Western blot analysis of phospho-RNA Pol II CTD serine 2 and serine 5 levels in zebrafish embryos after 24 h treatment with KI-DX-014 and reference CDK9 inhibitors.

### KI-DX-014 inhibits P-TEFb-dependent phosphorylation of RNA polymerase II CTD in zebrafish embryos

Since KI-DX-014 inhibited the release of P-TEFb from the 7SK snRNP in vitro, we sought to investigate its molecular and physiological effects in vivo. The structure of KI-DX-014, which includes an amino acid moiety (Fig. 1d), suggests poor membrane permeability (CLogP: 0.75). To overcome this limitation, we injected KI-DX-014 into 1-cell stage zebrafish embryos (Fig. 5c), a well-established model system for studying both P-TEFb and DDX21 during development. Injection of KI-DX-014 led to dose-dependent larval malformations, including curved body phenotypes and defects in facial development. Similar effects were observed in zebrafish embryos treated with the CDK9 inhibitor Flavopiridol^55,56^ (Fig. 5d). Notably, no substantial differences in survival rates were observed between the compound-treated groups and the DMSO control at 24 hours post-fertilization (24 hpf) (Supplementary Table 1). These results indicate that the compounds are non-toxic to embryos at the doses used in this study.

We also evaluated the phosphorylation of the Pol II CTD in the zebrafish embryos at 24 hpf. Both KI-DX-014 and the CDK9 inhibitor significantly reduced the phosphorylation of Ser2 and Ser5, established targets of activated P-TEFb^57^, in a dose-dependent manner (Fig. 5e, Supplementary Fig. 6). Based on these findings, we conclude that KI-DX-014 suppresses DDX21-dependent phosphorylation of the RNA polymerase II CTD by inhibiting P-TEFb release, leading to developmental defects in zebrafish embryos.

## Discussion

Here, we present the discovery of a small molecule probe that inhibits the binding of DDX21 to RNA and modulates functions that are dependent on RNA binding. KI-DX-001, discovered through SMM screening using lysate-based conditions, inhibits the binding of the stem-loop of 7SK, the endogenous substrate RNA of DDX21. Through SAR studies with KI-DX-001 analogues, we found an advanced probe, KI-DX-014, which directly binds to DDX21 confirmed by MST. Furthermore, a detailed analysis of RNA binding of 7SK using purified DDX21 domains revealed the DDX21 CTD to be the binding site of 7SK RNA. Additionally, KI-DX-014 competitively inhibits RNA binding at the DDX21 CTD. These results suggest that the DDX21 CTD is putative the binding site of KI-DX-014. ATPase activity, one of the RNA binding-dependent functions of DDX21, is allosterically regulated by RNA binding inhibition by KI-DX-014. Moreover, LLPS of RNA-bound DDX21 is also modulated by RNA binding inhibition of KI-DX-014. These findings that the SMM lysate screening approach is useful for RNA-binding protein targets with context-dependent functions utilizing endogenous substrates or with multifaceted functions, under conditions that more closely mimic a physiological environment, relative to binding screens involving purified proteins.

RNA binding proteins form biomolecular condensates through protein-protein or protein-RNA interactions and serve as sites for various biological processes. 7SK RNA-bound DDX21 is recruited to the promoter region of Pol II to promote the release of P-TEFb and to form transcriptional condensates that facilitate transcription^3^. In this study, we found that inhibition of RNA binding by KI-DX-014 can modulate the morphology of the phase separation formed by DDX21-RNA interactions. This suggests that RNA is a crucial structural component in the DDX21 phase separation. Furthermore, KI-DX-014 can be a useful chemical probe for investigating biological processes, which are facilitated by the DDX21 biomolecular condensates. In addition, the targeting of protein-RNA interactions could potentially provide a new strategy for the control of biomolecular condensates.

DDX21 has been shown to have the ability to release P-TEFb. P-TEFb facilitates Pol II elongation and promoter proximal escape^3^. In this study, we found that KI-DX-014 inhibited P-TEFb release via the inhibition of RNA binding. In zebrafish models, general transcriptional inhibition via enzymatic inhibition of CDK9 induces a developmental phenotype. Similarly, KI-DX-014 exhibited a developmental phenotype, and we found that it regulates the same pathway by analyzing the phosphorylation of Pol II CTD. KI-DX-014 can serve as a valuable chemical tool to investigate the role of DDX21 in the DDX21-P-TEFb-general transcription axis, as well as in DDX21-mediated tissue differentiation, cancer, and other diseases.

This study found that the chemical probe KI-DX-014 inhibited the binding of DDX21 RNA, but KI-DX-014 was not validated in living cells due to poor cell membrane permeability. Additionally, the precise mode of binding has not yet been determined. We believe that additional chemical optimization can improve the activity and physicochemical properties for analysis in living cell systems. Structural biology analysis, such as X-ray crystallography, can also help to define the precise binding mode and enable further medicinal chemistry. Furthermore, we believe that more selective RNA binding inhibitors will enable us to better understand DDX21-specific biology. Moreover, the regulation of general transcription factors by RNA binding inhibition of DDX21 can be further elucidated through global RNA expression analysis, which will provide a more detailed mechanistic characterization. Given our findings presented here, KI-DX-014 can serve as a valuable research tool for further investigations of the role of DDX21 in health and disease.

## Materials and Methods

### Cell culture

HEK293 cells were obtained from ATCC and grown under standard conditions in DMEM supplemented with 10% fatal bovine serum and Penicillin-Streptomycin-Glutamine (Gibco). To generate HEK cells expressing Halo-DDX21, we generated PiggyBac transposon vector (System Biosciences) containing tetracycline-inducible Halo-DDX21. The full-length human DDX21 CDS was cloned into a pHTN vector (Promega). Subsequently, the Halo-DDX21 fragment was amplified by PCR and cloned into tetracycline-inducible PiggyBac vector. The Halo-GFP expression vector was constructed by amplifying the GFP-coding sequence from the complementary DNA and inserting it into the tetracycline-inducible PiggyBac vector. To generate Halo-DDX21 fragments (HC+CTD, and CTD domains), gene fragments encoding the human DDX21 domain-coding region were synthesized (CTD: Invitrogen, HC+CTD: IDT). These synthesized gene fragments were then cloned into a PiggyBac vector that was both Halo-expressing and tetracycline-inducible. HEK293 cells were co-transfected with PiggyBac vector and PiggyBac transposase plasmid using Lipofectamine 2000 (Invitrogen). At 72 h post-transfection, stably integrated cells were selected using puromycin (Gibco) for two days. For transgene expression, the cells were cultured with the culture medium supplemented with doxycycline (Sigma) at 2 µg/mL for 48 h.

### SMM screening

Small molecule microarrays were manufactured as previously described^32^. Approximately 10,000 printed features were present on each SMM slide, consisting of 5,000 unique compounds printed in duplicate. 50K compound library and 65K compound library; 115,805 compounds in total were screened. HEK293 cells expressing Halo-DDX21 and Halo-GFP were cultured in growth medium (DMEM supplemented with 10% fetal bovine serum and Penicillin-Streptomycin-Glutamine (Gibco)). Doxycycline (2 µg/mL) was added to the medium to induce Halo-DDX21 and Halo-GFP expression. After 48 h, the cells were suspended in hypotonic lysis buffer (10 mM Tris-HCl, 10 mM NaCl, 5 mM MgCl_2_, pH 7.5) supplemented with 50X protease inhibitor cocktail (Promega) and incubated on ice for 15 min. 10% Triton X-100 was added to the suspensions at a final concentration of 0.2% and inverted five times. Then, the suspensions were centrifuged (5 min, 1,200 × g, 4 °C), and the supernatants were discarded to obtain nuclear pellets. The nuclear pellets were resuspended in mammalian lysis buffer (Promega) supplemented with a 50X protease inhibitor cocktail (Promega). To digest genomic DNA, RQ1 RNase-free DNase (Promega) was added to the lysates and incubated for 15 min at room temperature. The lysates were cleared by centrifugation (20 min, 16,000 × g, 4 °C). The concentration of Halo-fused protein in the lysate was determined by running HaloTag® TMR ligand (Promega)-treated lysates and a HaloTag® standard protein (Promega) standard curve on a Tris-Glycine gel. The gels were visualized using the ChemiDoc MP imaging system (Bio-Rad).

For SMM screening, the Halo-fused protein in the lysates was labeled with HaloTag® Alexa Fluor 660 ligand (Promega) for 30 min and diluted with binding buffer (50 mM HEPES, 150 mM NaCl, 0.005% IGEPAL CA-630, 0.1X Blocker FL (Thermo Fisher Scientific)) to a final concentration of 10 nM. The diluted lysate (2.5 mL) was applied to each SMM slide and incubated for 1 hour. Subsequently, the slides were subjected to three successive washes with TBS-T followed by a brief rinse in TBS and ultrapure water. The slides were then dried by centrifugation, after which they were immediately scanned using a SureScan Microarray Scanner (Agilent).

### SMM Data Analysis

The analysis of the scanned slide images was conducted by using GenePix Pro software (Molecular Devices) for alignment with the corresponding GenePix Array List (GAL) file, and the resulting GPR files were processed using data analysis workflow built by Pipeline Pilot (BIOVIA) and KNIME Analytics Platform. The robust Z-score for each feature was calculated as

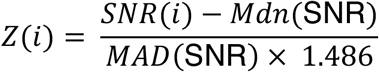

The signal-to-noise ratio for an individual feature is denoted by SNR(i). The median SNR value across all features in the subarray is represented by Mdn(SNR). MAD(SNR) indicates the maximum absolute deviation of SNR values for the entire set of features within the subarray. Assay positives were identified using the following criteria of each compound: robust Z-score > 3 in 4 replicates, SNR > 1.3 in 4 replicates for 65K library set. Halo-DDX21 slides were compared to slides for a Halo-GFP control protein. Assay positives were verified by manual inspection of slide images.

### Recombinant Protein Purification

HEK293 cells expressing Halo-fused target protein were cultured in growth media (DMEM supplemented with 10% fatal bovine serum and Penicillin-Streptomycin-Glutamine (Gibco)). Doxycycline (2 µg/mL) was added to the medium to induce Halo-DDX21 and Halo-GFP expression. After 48 h, cells were lysed in mammalian lysis buffer (Promega) supplemented with 50X Protease Inhibitor Cocktail (Promega). RQ RNase-free DNase (Promega) and RNase A (NEB) were added to the lysate and incubate for 15 min at room temperature and then the lysate was centrifuged (10 min, 16,000 x g, 4 °C). Pre-washed Magne^TM^ HaloTag® Beads (Promega) was added to the lysate and incubated at 4 °C for overnight. The beads were subjected to three washes with wash buffer (50 mM Tris-HCl, 3 mM EDTA, 1 M NaCl, 0.1 mM DTT, 0.5% NP-40, 10% Glycerol, pH 7.5) and subsequently underwent three additional washes with digestion buffer (50 mM HEPES, 150 mM NaCl, 1 mM DTT, 0.005% IGEPAL CA-630, pH 7.5). The beads were transferred to Low protein binding microcentrifuge tube (Thermo Fisher Scientific) and then added HaloTEV protease (Promega) in digestion buffer. The beads were incubated for 90 minutes at 30 °C, 1000 rpm. After 90 minutes, the eluate was collected, and the beads were washed once with digestion buffer. The eluate and washed buffer were combined and concentrated using Amicon Ultra-0.5 30K molecular weight cutoff. To quantify the target protein, samples were subjected to Tris-Glycine gel electrophoresis alongside a BSA standard curve. The gel was then stained with SimplyBlue SafeStain (Invitrogen) to visualize the proteins. The gel was visualized using Azure imager (Azure Biosystems). The protein band intensity was analyzed using imageJ-Fiji software.

### Fluorescence Anisotropy Assay

50 µM 5’-FAM-labeled RNA was heated at 95 °C for 5 min in folding buffer (20 mM HEPES, 1 mM MgCl_2_, 100 mM NaCl, 1 mM DTT, 10% Glycerol, pH 7.5) and then placed on ice. For RNA probe binding experiments, proteins were incubated with 0.5 nM 5’-FAM-labeled RNA in FP buffer (20 mM Tris-acetate, 50 mM KCl, 2 mM MgCl2, 1 mM DTT, 0.01% IGEPAL CA-630, 1 mM AMPPCP (Sigma), pH 7.5) in black 384-well plates (Greiner, 781209) for 1 hour at room temperature. For the competition assay, proteins were pre-incubated with compounds (or 5 µM non-labeled 7SK-SL4 or DMSO) for 30 min at room temperature and subsequently incubated with 0.5 nM 5’-FAM-labeled RNA in FP buffer for 1 hour at room temperature. Fluorescence anisotropy was measured using Spark (Tecan).The K_d_ was determined by applying a monovalent reversible equilibrium binding model that accounts for ligand depletion, using the method previously described^58^.

### Electrophoretic Mobility Shift Assay (EMSA)

50 µM 5’-IRDye800CW-7SK-SL4 (IDT) was heated at 95 °C for 5 min in folding buffer (20 mM HEPES, 1 mM MgCl_2_, 100 mM NaCl, 1 mM DTT, 10% Glycerol, pH 7.5). Binding reactions containing 15 nM purified full-length DDX21 (or RNase-free water), ligand (or DMSO (Sigma)), 1 nM 5’-IRDye800CW-7SK-SL4 (IDT) in binding buffer (20 mM Tris-acetate, 50 mM KCl, 2 mM MgCl_2_, 1 mM AMPPCP (Sigma), pH 7.5) were prepared in PCR tubes and incubated at room temperature for 1 hour. Each reaction was supplemented with 10X Orange loading dye (Li-Cor) prior to being loaded onto a 5% Mini-Protean TBE gel (Bio-Rad). The gel was run at 100 V for 30 minutes in 1X TBE before applying the samples. Then, the samples underwent electrophoresis at 100 V for 40 minutes at room temperature. Subsequently, the gels were imaged using an Odessey CLx imager (Li-Cor).

### Microscale Thermophoresis (MST)

The His-Tag Labeling Kit RED-tris-NTA 2nd Generation (NanoTemper Technologies, MO-L018) was employed to label purified DDX21 HC+CTD. The reaction was conducted in a reaction mixture comprising 100 nM of the labeled protein in a binding buffer (20 mM Tris-HCl, 50 mM KCl, 2 mM MgCl_2_, 1 mM DTT, 0.05% IGEPAL CA-630, pH 7.5) for 30 min at room temperature in the dark. The labeled protein was diluted to 10 nM with binding buffer containing Blocker FL (Thermo Fisher Scientific) and centrifuged (5 min, 13,000 × g, 4 °C) to remove protein aggregates. The labeled protein was mixed with KI-DX-014 in equivalent ratios and allowed to equilibrate for 15 minutes at room temperature before being transferred to Monolith Capillaries (NanoTemper Technologies, MO-K022). The MST experiment was carried out at 25 °C, employing a Monolith NT.115 pico (NanoTemper Technologies) set to 40% excitation power and Medium MST power using. The dissociation constant (Kd) was calculated by evaluating data from four discrete pipetted measurements, employing a Kd Model through MO. Affinity Analysis software (NanoTemper Technologies). The data were normalized by the fraction bound and are indicated in the figure.

### *In Vitro* ATPase Activity Assay

The ATPase activity of DDX21 was evaluated utilizing the ADP-Glo assay (Promega) in accordance with the manufacturer’s protocol. The reactions were performed with 100 nM of full-length DDX21 and RNA substrate (either 100 nM 7SK-SL4 or 1 µM U15) in a reaction buffer (40 mM Tris-acetate, 20 mM MgCl_2_, and 0.1 mg/mL BSA at pH 7.5). The reactions were performed in white 384-well plates (BrandTech Scientific, 781621). The resultant luminescence was measured using INFINITE M200 PRO (Tecan) instrument.

### Helicase RNA Unwinding Assay

The evaluation of DDX21-mediated RNA unwinding was conducted in accordance with a previously established protocol^59^ with minor modifications. The reaction mixture containing 200 nM full-length DDX21, 30 nM dsRNA probes (IDT), 300 nM complementary DNA (IDT), and KI-DX-014 (or DMSO) in an unwinding buffer (20 mM Tris-acetate, 2 mM Magnesium acetate, 100 mM KCl, 0.2 mM DTT, and 0.4 U/µL RiboLock RNase Inhibitor (Invitrogen)) was placed in black 384 well plates (Greiner, 781209) and incubated with 1 mM Mg-ATP at 37 °C overnight. The resulting fluorescence intensity was measured using INFINITE M200 PRO (Tecan) instrument.

### Droplet Formation Assay

The reactions containing 10 µM full length DDX21, KI-DX-014 or DMSO or 0.4 mg/mL RNase A (New England Biolabs) and 2 µM 5’-FAM-labeled RNA (7SK-SL4 or U15) in reaction buffer (20 mM Tris-HCl, 50 mM KCl, 2 mM MgCl_2_, 10% PEG8000) were incubated for 30 min at room temperature. 7 µL of the reaction was dropped onto glass slide (Fisher Scientific) then covered by coverslip (Fisher Scientific). Fluorescence image was obtained using EVOS system (Thermo Fisher Scientific). Average droplet size per image was calculated using CellProfiler (v 4.2.6). Significance was determined by one-way ANOVA test using Prism 10 (v 10.2.3)

### Immunoblotting

For immunoblotting studies, samples were loaded onto 4-12% or 4-20% Novex Tris-Glycine gel (Invitrogen) and subjected to electrophoresis at 250 V for 30 min. Proteins were subsequently transferred onto a PVDF membrane (Merck) at 100 V for 1 h at room temperature or 30 V overnight at 4 °C and then stained utilizing Revert 700 Total Protein Stain Kits (Li-Cor) for total protein normalization. The membrane was blocked with EveryBlot (Bio-Rad) at room temperature for 10 min. The membrane was then incubated with primary antibody at 4 °C overnight or at room temperature for 2 hours. Membranes were washed with PBS-T and subsequently incubated with secondary antibody at room temperature for 1 hour, followed by washing with PBS-T. Signals were detected via chemiluminescence on an Azure imager (Azure Biosystems) utilizing SuperSignal West Femto Maximum Sensitivity Substrate (Thermo Fisher Scientific). The protein band intensity was analyzed using imageJ-Fiji software.

The antibodies used in this study include CDK9 (C12F7) Rabbit mAb (Cell Signaling, 2316, 1:1000), Anti-HEXIM1 antibody (Abcam, ab25388, 1:1000), Purified anti-RNA Polymerase II RPB1 Antibody (H5) (BioLegend, 920204, 1:1000), Anti-RNA polymerase II CTD repeat YSPTSPS (phospho S5) antibody (Abcam, ab5131, 1:1000), Goat anti-Mouse IgG (H+L) Poly-HRP Secondary Antibody, HRP (Invitrogen, 32230, 1:5000), Goat anti-Rabbit IgG (H+L) Poly-HRP Secondary Antibody, HRP (Invitrogen, 32260, 1:5000), Peroxidase IgG Fraction Monoclonal Mouse Anti-Rabbit IgG, light chain specific (Jackson ImmunoResearch, 211-032-171, 1:5000)

### P-TEFb Release Assay

The P-TEFb release assay was performed in accordance with a previously established protocol^3^ with minor modifications. 1.25 µg of anti-HEXIM1 antibody (Abcam, ab25388) was bound to Dynabeads protein G (Invitrogen) and incubated with 1 mg of HeLa cell nuclear extracts at 4 °C for overnight to immobilize the inactive 7SK snRNP complex. The resulting immunocomplexes were incubated with 30 µg of RNase A (New England Biolabs) or 3 µg of purified full-length DDX21 with increasing amounts of KI-DX-014 and 1mM AMPPCP (Sigma) or 1 mM ATP for 1 min, 1350 rpm/ 3 min, OFF, 30 cycles at 4 °C. The supernatants were separated using a magnetic separator and supplemented with 4X LDS sample buffer, followed by heating for 5 minutes at 95°C. The remaining beads were subjected to heating in 1X LDS sample buffer for 10 minutes at 70°C. Subsequently, the samples were analyzed by Western blotting.

### Zebrafish Handling

Zebrafish lines were housed in AAALAC-approved facilities and maintained according to protocols approved by the Massachusetts Institute of Technology Committee on Animal Care. Experiments were performed in the AB/Tübingen (TAB5/14) genetic background. Wild-type zebrafish were used in all experiments. For drug microinjection experiments, 1-or 2-cell-stage zebrafish were injected with 1 nL of KI-DX-014 (2.5, 5, and 10 mM in 50% (v/v) DMSO/water) or 50% (v/v) DMSO/water as a control and incubated at 28 °C for 24 h. For drug treatment, 1-or 2-cell-stage zebrafish were incubated in 3 mL of flavopiridol (0.25, 0.5, and 1 µM in 0.5% (v/v) DMSO/water), or 0.5% (v/v) DMSO/water as control and incubated at 28 °C for 24 h.

### Total Protein Preparation from Zebrafish Embryos

The embryo chorion was surgically removed using fine tweezers and transferred into a 1.5 mL microcentrifuge tube. Dechorionated embryos were suspended in ice-cold PBS, pipetted up and down to remove the yolk sacs, and washed once with ice-cold PBS. The dechorionated and de-yolked larvae were centrifuged for 30 s at 20,000 × g at room temperature and the supernatant was removed. RIPA buffer supplemented with protease inhibitor cocktail, PSMF, and PhosSTOP (Roche) was added and homogenized with a microfuge pestle for 45 s and then sonicated for 30 s ON/ 30 s OFF for 5 cycles using Bioruptor (Diagenode). The homogenates were subjected to centrifugation (10 min, 16,000 × g, 4 °C). The supernatant was subsequently collected and transferred to a new 1.5 mL microcentrifuge tube. The protein concentration of the samples was determined using the Bradford assay. An equivalent quantity of total protein was transferred into a new microcentrifuge tube, and 4X LDS sample buffer (Thermo Fisher Scientific) was added. The samples were then heated at 95 °C for 5 min and analyzed by western blotting.

## Acknowledgements

We thank A. Mohamed from the Vos lab at MIT for providing 5’-FAM labeled 7SK-SL1 and SL4. We also thank the Center for Macromolecular Interactions at Harvard Medical School for access to MST instrumentation (RRID:SCR_018270) and the Koch Institute Zebrafish Core Facility. This work was supported by Ono Pharmaceutical Co., Ltd., and Koch Institute Support (core) Grant 5P30-CA014051 from the National Cancer Institute.

---

**Supplementary Figure 1:**
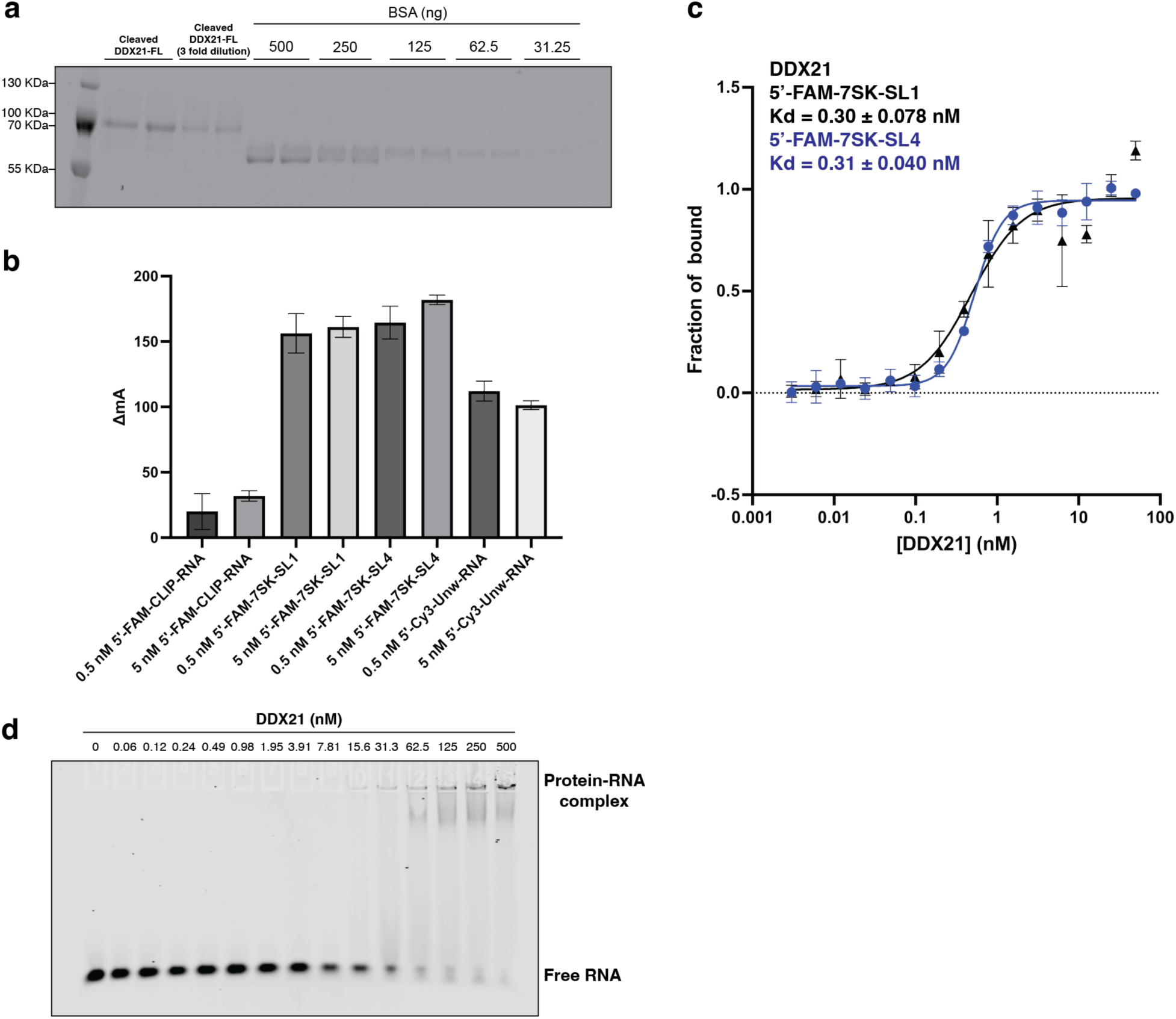
Purification of DDX21-FL and development of DDX21-RNA binding assays. **a**. Coomassie stain of SDS-PAGE gel for quantification purified DDX21-FL. **b**, Development of the fluorescence anisotropy assay using DDX21-FL and RNA probe; 5’-FAM-CLIP-RNA, 5’-FAM-7SK-SL1, 5’-FAM-7SK-SL4 and 5’-Cy3-Unw-RNA. ΔmA (y-axis) was calculated as the difference between the mA value for each probe and the value for the condition without protein addition. Data are shown as mean values ± s.d. for three technical replicates. **c**, Fluorescence anisotropy assay for competition of 5’-FAM labeled 7SK-SL4 RNA for binding to DDX21-FL using KI-DX-014. Data are shown as mean values ± s.d. for three technical replicates. **d**. Electrophoretic mobility shift assay (EMSA) of DDX21-FL for confirmation of RNA binding using IRDye800CW-labeled 7SK-SL4.

**Supplementary Figure 2:**
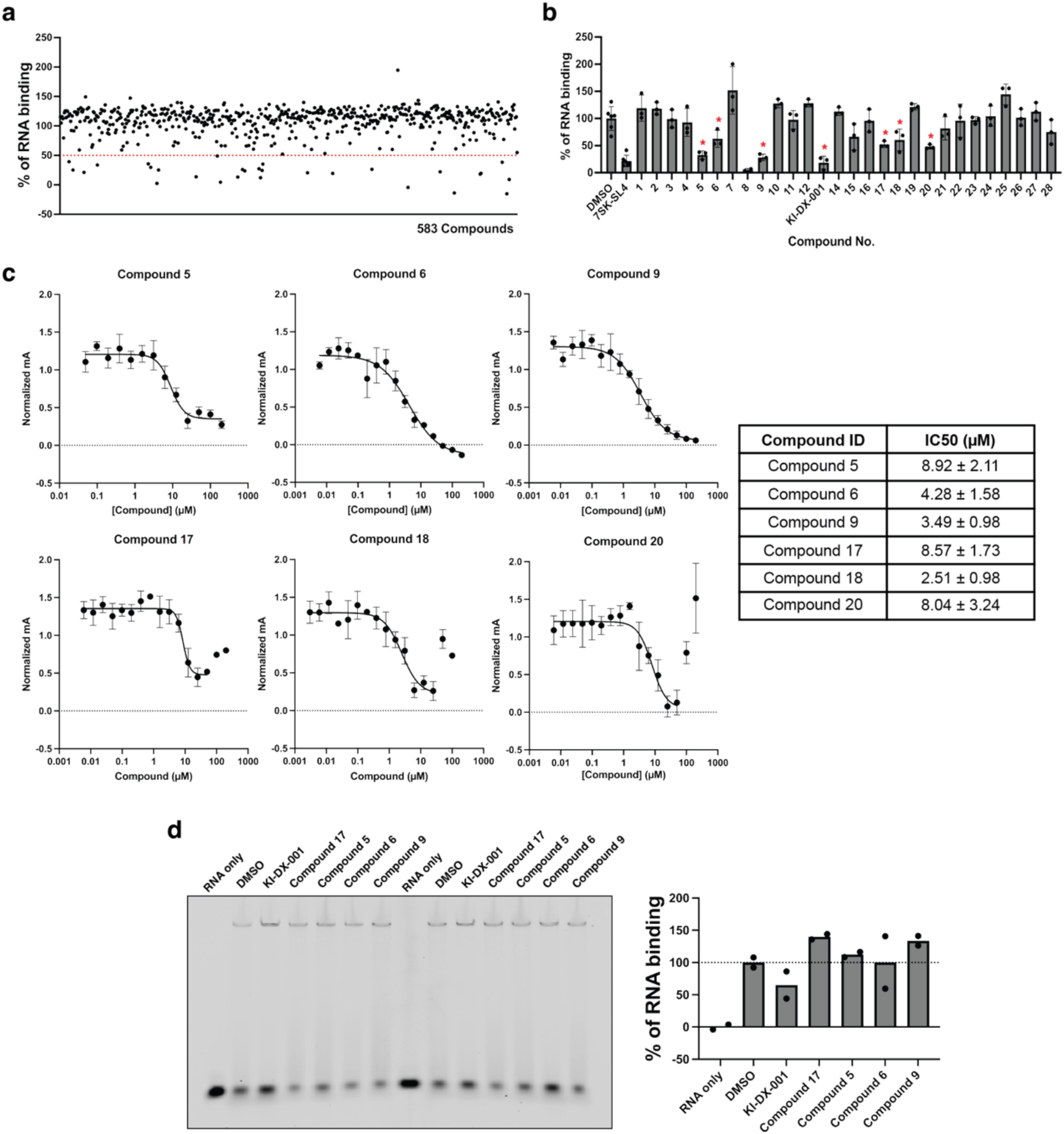
Discovery and *in vitro* biological evaluation of KI-DX-001. **a**. Secondary fluorescence anisotropy assay screening for primary SMM hits using cherry picked compounds from library stocks at 20 µM. A threshold of 50% inhibition is shown red dashed line. **b**. Confirmation assay for secondary fluorescence anisotropy assay screening using re-purchased compounds in triplicates at 20 µM. Compounds which marked with asterisk were showed over 35% inhibition in triplicates. **c**, Fluorescence anisotropy assay for competition of 5’-FAM labeled 7SK-SL4 RNA for binding to DDX21-FL. Data are shown as mean values ± s.d. for three technical replicates. **d**. EMSA of DDX21-FL-RNA binding inhibition using IRDye800CW labeled 7SK-SL4 after treatment with validated compounds from secondary screening; Free RNA band intensities were quantified using Fiji 2.14.0.

**Supplementary Figure 3:**
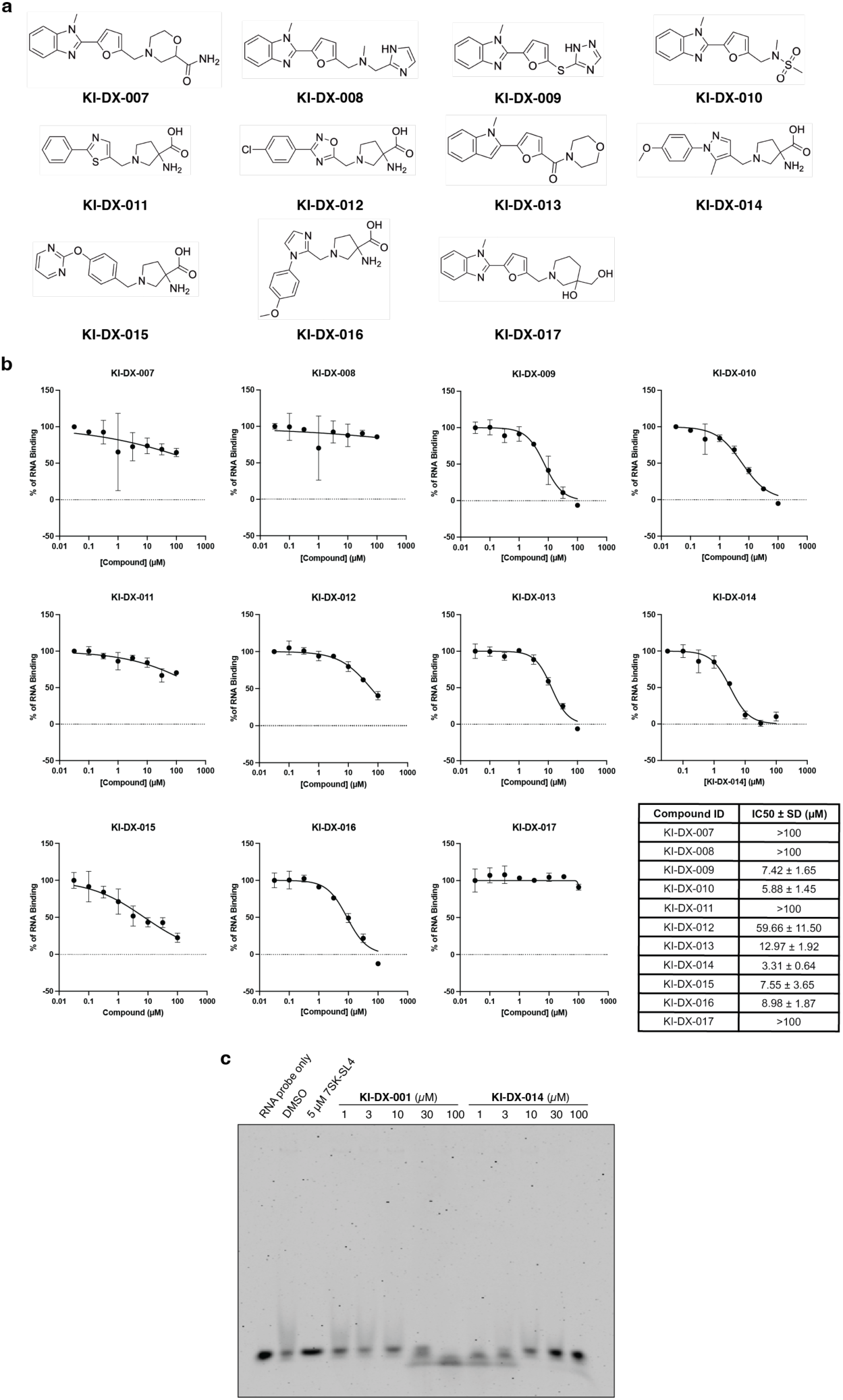
Hit expansion to discover advanced hit probe, KI-DX-014 **a**. Chemical structures of KI-DX-001 analogues. **b**, Fluorescence anisotropy assay for competition of 5’-FAM labeled 7SK-SL4 RNA for binding to DDX21-FL using KI-DX-001 analogues. Data are shown as mean values ± s.d. for three technical replicates. **c**, EMSA of DDX21-FL-RNA binding inhibition using 5’-IRDye800CW labeled 7SK-SL4 after treatment with KI-DX-001 and KI-DX-014.

**Supplementary Figure 4:**
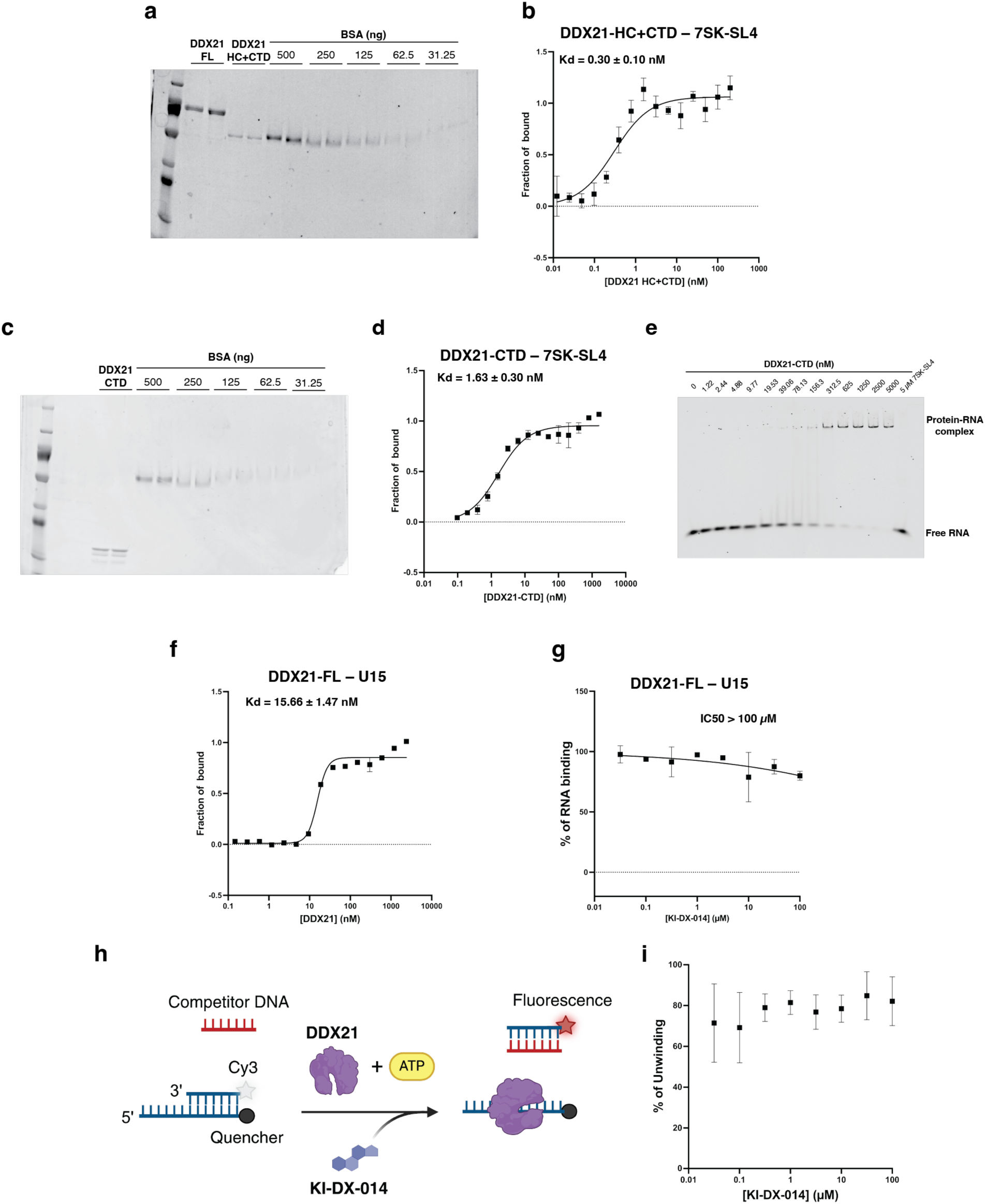
Characterization of *in vitro* biochemical activity of DDX21 full length and truncated constructs. **a**. Coomassie stain of SDS-PAGE gel for quantification of purified DDX21-HC+CTD. **b**. Fluorescence anisotropy assay for 5’-FAM labeled 7SK-SL4 RNA binding to DDX21-HC+CTD. Data are shown as mean values ± s.d. for three technical replicates. **c**. Coomassie stain of SDS-PAGE gel for quantification of purified DDX21-CTD. **d**. Fluorescence anisotropy assay for 5’-FAM labeled 7SK-SL4 RNA binding to DDX21-CTD. Data are shown as mean values ± s.d. for three technical replicates. **e**. EMSA of DDX21-CTD-RNA binding using 5’-IRDye800CW labeled 7SK-SL4. **f**. Dose-response curves for the fluorescence anisotropy assay of DDX21-CTD-RNA binding using 5’-FAM labeled U15. (n = 3 technical replicates, error bars represent means ± s.d.) **g**. Fluorescence anisotropy assay for competition of 5’-FAM labeled U15 RNA for binding to DDX21-U15. Data are shown as mean values ± s.d. for three technical replicates. **h**. Schematic of helicase unwinding assay. Created with <u>BioRender.com</u>. **i**. The helicase activity of DDX21-FL in the presence of KI-DX-014. (n = 3 technical replicates, error bars represent means ± s.d.)

**Supplementary Figure 5:**
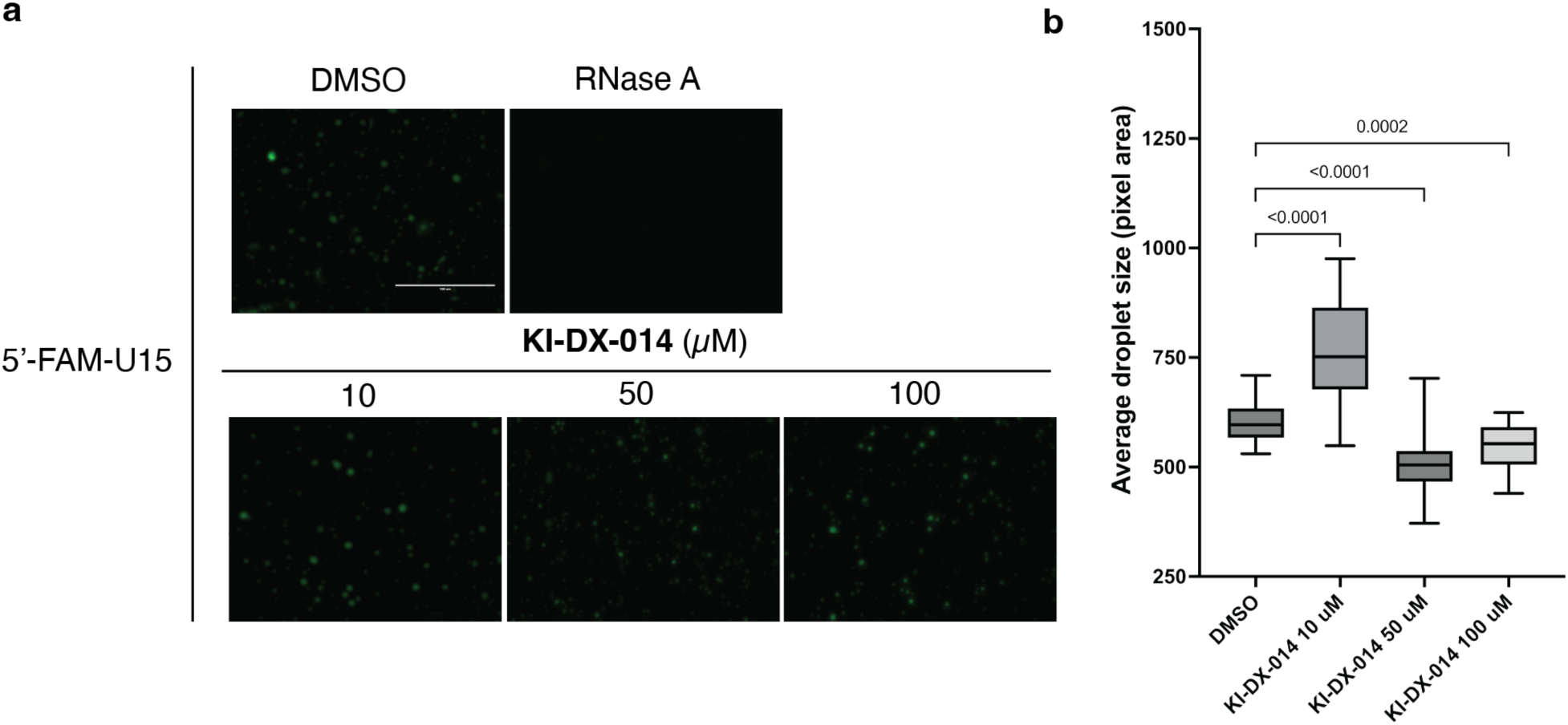
Characterization of RNA binding-dependent DDX21 activities using U15 as an RNA substrate **a.** Concentration dependent effects of KI-DX-014 on the morphological properties of DDX21 condensates, showing representative fluorescence images with 5’-FAM labeled U15 in the presence of KI-DX-014 or RNase A in 10% PEG8000; scale bar, 100 µm. **b**. Quantification of the average size of the droplets. Data are shown using box–whisker plots and error bars represents s.e.m.. *P* values were determined by one-way ANOVA with Dunnett’s test for multiple comparison.

**Supplementary Fig. 6:**
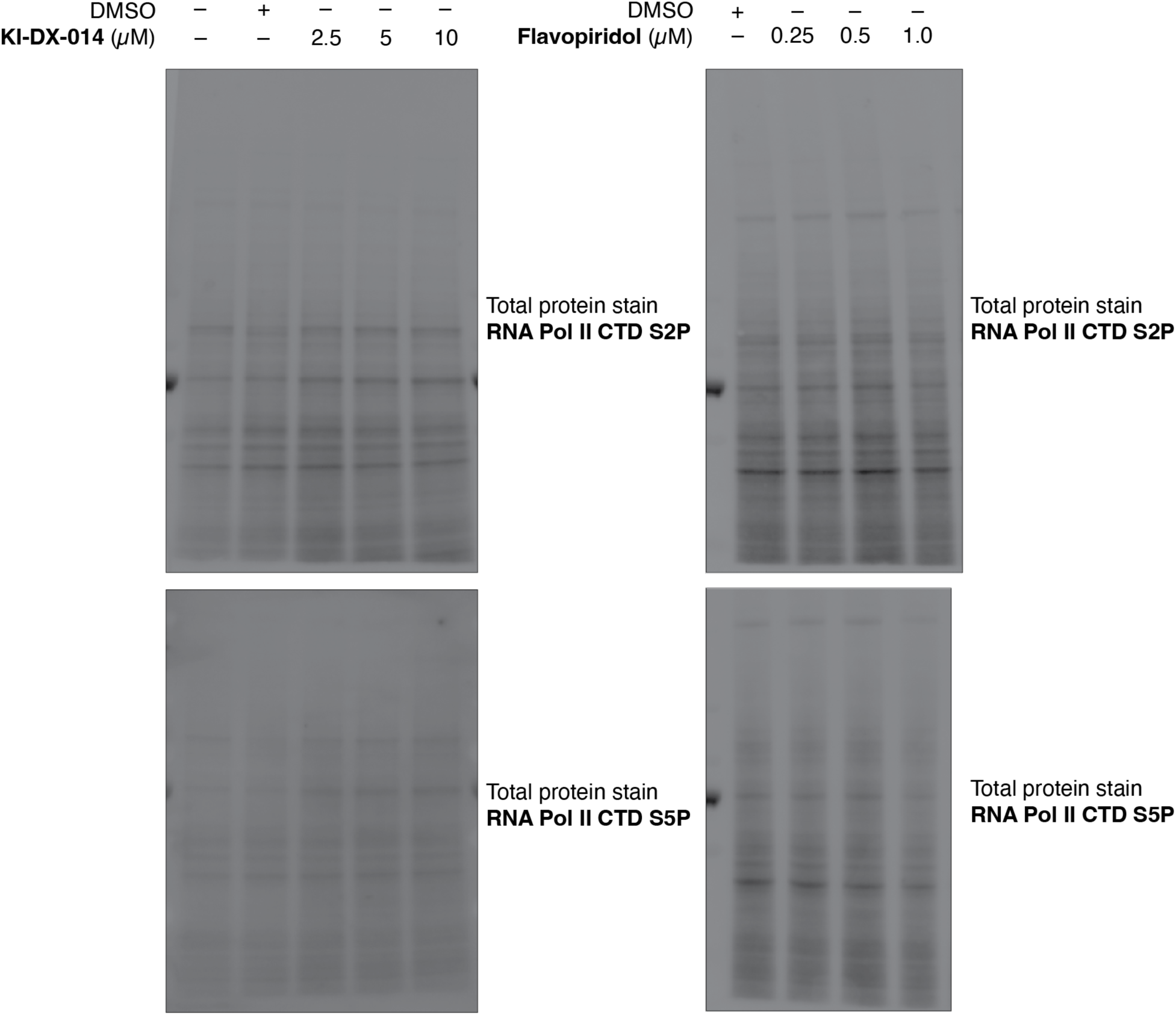
Immunoblotting of total protein of zebrafish embryos after treatment with KI-DX-014 and reference CDK9 inhibitor for 24 hours.

**Supplementary Table 1.**
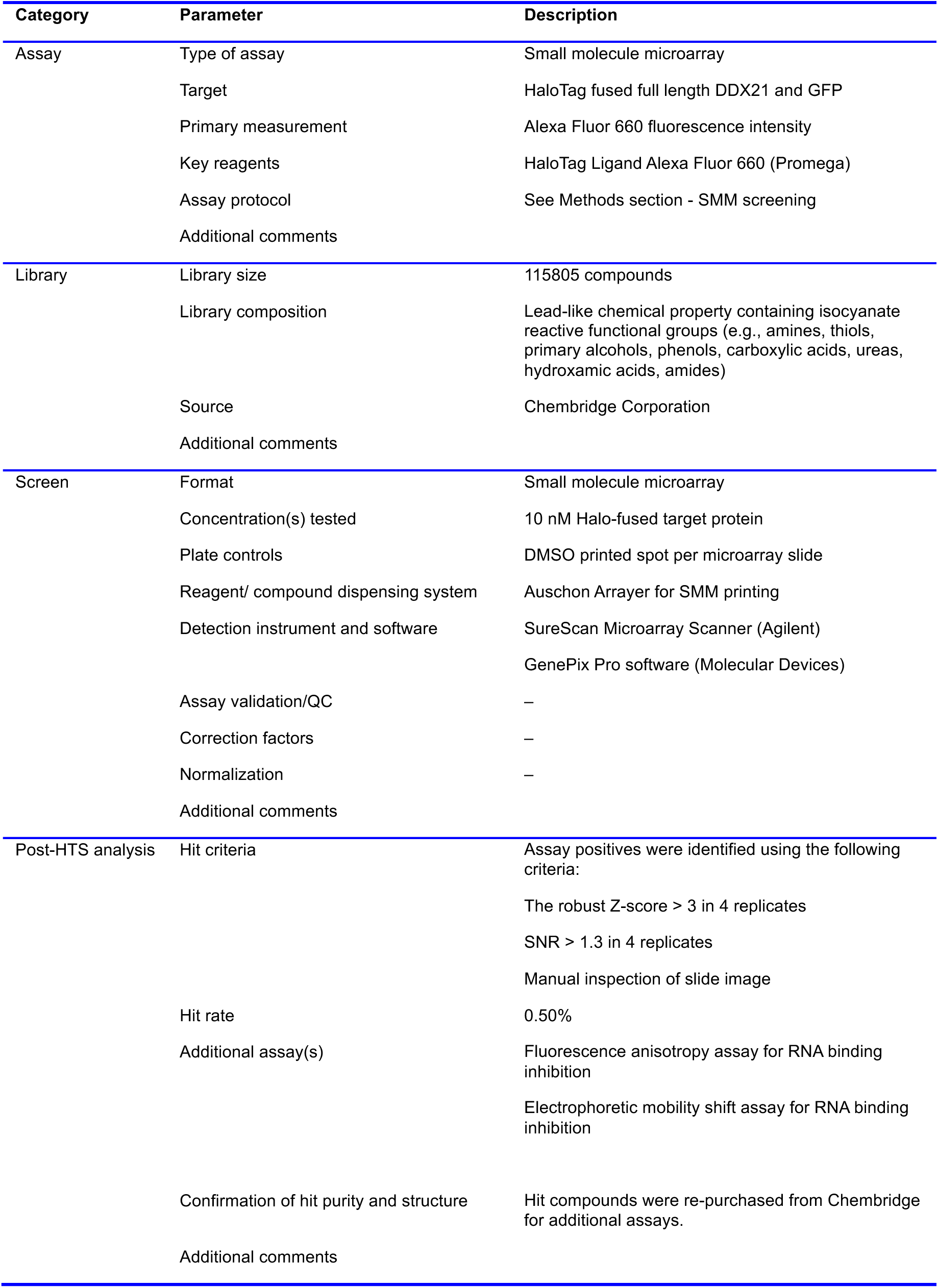
Small molecule screening data.

**Supplementary Table 2.** Survival rate of zebrafish embryos after treatment with KI-DX-014 and reference CDK9 inhibitor for 24 hours.

| Conditions | %Survival |
| --- | --- |
| DMSO (Injection) | 73.2% |
| 2.5 μM KI-DX-014 | 69.2% |
| 5 μM KI-DX-014 | 60.6% |
| 10 μM KI-DX-014 | 71.1% |
| DMSO (Treatment) | 66.7% |
| 1 μM Flavopiridol | 62.3% |

## Reference

1 Flores-Rozas, H. & Hurwitz, J. Characterization of a new RNA helicase from nuclear extracts of HeLa cells which translocates in the 5‘ to 3‘ direction. J. Biol. Chem. 268, 21372–21383 (1993). 10.1016/s0021-9258(19)36933-9

2 Hammond, J. A. et al. A Survey of DDX21 Activity During Rev/RRE Complex Formation. J. Mol. Biol. 430, 537–553 (2018). 10.1016/j.jmb.2017.06.023

3 Calo, E. et al. RNA helicase DDX21 coordinates transcription and ribosomal RNA processing. Nature 518, 249–253 (2014). 10.1038/nature13923

4 Henning, D., So, R. B., Jin, R., Lau, L. F. & Valdez, B. C. Silencing of RNA helicase II/Gualpha inhibits mammalian ribosomal RNA production. J. Biol. Chem. 278, 52307–52314 (2003). 10.1074/jbc.M310846200

5 Miao, W. et al. Glucose dissociates DDX21 dimers to regulate mRNA splicing and tissue differentiation. Cell 186, 80–97 (2023). 10.1016/j.cell.2022.12.004

6 Johansson, J. A. et al. PRL3-DDX21 Transcriptional Control of Endolysosomal Genes Restricts Melanocyte Stem Cell Differentiation. Dev. Cell 54, 317–332 (2020). 10.1016/j.devcel.2020.06.013

7 Chen, Z. et al. Structural Basis of Human Helicase DDX21 in RNA Binding, Unwinding, and Antiviral Signal Activation. Adv. Sci. 7, 2000532 (2020). 10.1002/advs.202000532

8 Marcaida, M. J. et al. The Human RNA Helicase DDX21 Presents a Dimerization Interface Necessary for Helicase Activity. iScience 23 (2020). 10.1016/j.isci.2020.101811

9 Peterlin, B. M., Brogie, J. E. & Price, D. H. 7SK snRNA: a noncoding RNA that plays a major role in regulating eukaryotic transcription. Wiley Interdiscip. Rev. RNA. 3, 92–103 (2011). 10.1002/wrna.106

10 Fujinaga, K., Huang, F. & Peterlin, B. M. P-TEFb: The master regulator of transcription elongation. Mol. Cell 83, 393–403 (2023). 10.1016/j.molcel.2022.12.006

11 Lin, Y., Protter, David S. W., Rosen, Michael K. & Parker, R. Formation and Maturation of Phase-Separated Liquid Droplets by RNA-Binding Proteins. Mol. Cell 60, 208–219 (2015). 10.1016/j.molcel.2015.08.018

12 Aryan, F. et al. Nucleolus activity-dependent recruitment and biomolecular condensation by pH sensing. Mol. Cell 83, 4413–4423 (2023). 10.1016/j.molcel.2023.10.031

13 Wu, M. et al. lncRNA SLERT controls phase separation of FC/DFCs to facilitate Pol I transcription. Science 373, 547–555 (2021). 10.1126/science.abf6582

14 Xing, Y. H. et al. SLERT Regulates DDX21 Rings Associated with Pol I Transcription. Cell 169, 664–678 (2017). 10.1016/j.cell.2017.04.011

15 Gao, H. et al. Phase separation of DDX21 promotes colorectal cancer metastasis via MCM5-dependent EMT pathway. Oncogene 42, 1704–1715 (2023). 10.1038/s41388-023-02687-6

16 Bora, P. et al. DDX21 is a p38-MAPK-sensitive nucleolar protein necessary for mouse preimplantation embryo development and cell-fate specification. Open Biol. 11, 210092 (2021). 10.1098/rsob.210092

17 Calo, E. et al. Tissue-selective effects of nucleolar stress and rDNA damage in developmental disorders. Nature 554, 112–117 (2018). 10.1038/nature25449

18 Santoriello, C. et al. RNA helicase DDX21 mediates nucleotide stress responses in neural crest and melanoma cells. Nat. Cell Biol. 22, 372–379 (2020). 10.1038/s41556-020-0493-0

19 Song, C., Hotz-Wagenblatt, A., Voit, R. & Grummt, I. SIRT7 and the DEAD-box helicase DDX21 cooperate to resolve genomic R loops and safeguard genome stability. Genes Dev. 31, 1370–1381 (2017). 10.1101/gad.300624.117

20 Zhang, Y., Baysac, K. C., Yee, L.-F., Saporita, A. J. & Weber, J. D. Elevated DDX21 regulates c-Jun activity and rRNA processing in human breast cancers. Breast Cancer Res. 14, 499 (2014). 10.1186/s13058-014-0449-z

21 Cao, J. et al. DDX21 promotes gastric cancer proliferation by regulating cell cycle. Biochem. Biophys. Res. Commun. 505, 1189–1194 (2018). 10.1016/j.bbrc.2018.10.060

22 Licciardello, M. P. & Workman, P. The era of high-quality chemical probes. RSC Med. Chem. 13, 1446–1459 (2022). 10.1039/d2md00291d

23 D’Agostino, V. G. et al. Screening Approaches for Targeting Ribonucleoprotein Complexes: A New Dimension for Drug Discovery. SLAS Discov. 24, 314–331 (2019). 10.1177/2472555218818065

24 Julio, A. R. & Backus, K. M. New approaches to target RNA binding proteins. Curr. Opin. Chem. Biol. 62, 13–23 (2021). 10.1016/j.cbpa.2020.12.006

25 Garcia-Jove Navarro, M., et al. RNA is a critical element for the sizing and the composition of phase-separated RNA–protein condensates. Nat. Commun. 10 (2019). 10.1038/s41467-019-11241-6

26 Vries, T. d., et al. Specific protein-RNA interactions are mostly preserved in biomolecular condensates. Sci. Adv. 10, 1–12 (2024). 10.1126/sciadv.adm7435

27 Wadsworth, G. M. et al. RNA-driven phase transitions in biomolecular condensates. Mol. Cell 84, 3692–3705 (2024). 10.1016/j.molcel.2024.09.005

28 Han, T. W., Portz, B., Young, R. A., Boija, A. & Klein, I. A. RNA and condensates: Disease implications and therapeutic opportunities. Cell Chem. Biol. 31, 1593–1609 (2024). 10.1016/j.chembiol.2024.08.009

29 Wu, P. Inhibition of RNA-binding proteins with small molecules. Nat. Rev. Chem. 4, 441–458 (2020). 10.1038/s41570-020-0201-4

30 Naineni, S. K., Robert, F., Nagar, B. & Pelletier, J. Targeting DEAD-box RNA helicases: The emergence of molecular staples. Wiley Interdiscip. Rev. RNA. 14 (2022). 10.1002/wrna.1738

31 Bradner, J. E. et al. A Robust Small-Molecule Microarray Platform for Screening Cell Lysates. Chem. Biol. 13, 493–504 (2006). 10.1016/j.chembiol.2006.03.004

32 Casalena, D. E., Wassaf, D. & Koehler, A. N. Ligand Discovery Using Small-Molecule Microarrays. Methods Mol. Biol., 249–263 (2012). 10.1007/978-1-61779-364-6_17

33 Vegas, A. J., Fuller, J. H. & Koehler, A. N. Small-molecule microarrays as tools in ligand discovery. Chem. Soc. Rev. 37 (2008). 10.1039/b703568n

34 Pop, M. S. et al. A small molecule that binds and inhibits the ETV1 transcription factor oncoprotein. Mol. Cancer Ther. 13, 1492–1502 (2014). 10.1158/1535-7163.MCT-13-0689

35 Struntz, N. B. et al. Stabilization of the Max Homodimer with a Small Molecule Attenuates Myc-Driven Transcription. Cell Chem. Biol. 26, 711–723 (2019). 10.1016/j.chembiol.2019.02.009

36 Richters, A. et al. Modulating Androgen Receptor-Driven Transcription in Prostate Cancer with Selective CDK9 Inhibitors. Cell Chem. Biol. 28, 134–147 (2021). 10.1016/j.chembiol.2020.10.001

37 Los, G. V. et al. HaloTag: A Novel Protein Labeling Technology for Cell Imaging and Protein Analysis. ACS Chem. Biol. 3, 373–382 (2008). 10.1021/cb800025k

38 Noblin, D. J. et al. A HaloTag-Based Small Molecule Microarray Screening Methodology with Increased Sensitivity and Multiplex Capabilities. ACS Chem. Biol. 7, 2055–2063 (2012). 10.1021/cb300453k

39 Quinnell, S. P. et al. A Small-Molecule Inhibitor to the Cytokine Interleukin-4. ACS Chem. Biol. 15, 2649–2654 (2020). 10.1021/acschembio.0c00615

40 Wang, S. et al. Current understanding of the role of DDX21 in orchestrating gene expression in health and diseases. Life Sci. 349, 122716 (2024). 10.1016/j.lfs.2024.122716

41 Irwin, J. J. et al. ZINC20—A Free Ultralarge-Scale Chemical Database for Ligand Discovery. J. Chem. Inf. Model. 60, 6065–6073 (2020). 10.1021/acs.jcim.0c00675

42 Jerabek-Willemsen, M. et al. MicroScale Thermophoresis: Interaction analysis and beyond. J. Mol. Struct. 1077, 101–113 (2014). 10.1016/j.molstruc.2014.03.009

43 McRae, E. K. S. et al. Human DDX21 binds and unwinds RNA guanine quadruplexes. Nucleic Acids Res. 45, 6656–6668 (2017). 10.1093/nar/gkx380

44 Valdez, B. C., Henning, D., Perumal, K. & Busch, H. RNA-unwinding and RNA-folding activities of RNA helicase II/Gu--two activities in separate domains of the same protein. Eur. J. Biochem. 250, 800–807 (1997). 10.1111/j.1432-1033.1997.00800.x

45 Zegzouti, H., Zdanovskaia, M., Hsiao, K. & Goueli, S. A. ADP-Glo: A Bioluminescent and homogeneous ADP monitoring assay for kinases. Assay Drug Dev. Technol. 7, 560–572 (2009). 10.1089/adt.2009.0222

46 Hondele, M. et al. DEAD-box ATPases are global regulators of phase-separated organelles. Nature 573, 144–148 (2019). 10.1038/s41586-019-1502-y

47 Wei, P., Garber, M. E., Fang, S.-M., Fischer, W. H. & Jones, K. A. A Novel CDK9-Associated C-Type Cyclin Interacts Directly with HIV-1 Tat and Mediates Its High-Affinity, Loop-Specific Binding to TAR RNA. Cell 92, 451–462 (1998). 10.1016/S0092-8674(00)80939-3

48 Peng, J., Zhu, Y., Milton, J. T. & Price, D. H. Identification of multiple cyclin subunits of human P-TEFb. Genes Dev. 12, 755–762 (1998). 10.1101/gad.12.5.755

49 Adelman, K. & Lis, J. T. Promoter-proximal pausing of RNA polymerase II: emerging roles in metazoans. Nat. Rev. Genet. 13, 720–731 (2012). 10.1038/nrg3293

50 Jang, M. K. et al. The bromodomain protein Brd4 is a positive regulatory component of P-TEFb and stimulates RNA polymerase II-dependent transcription. Mol. Cell 19, 523–534 (2005). 10.1016/j.molcel.2005.06.027

51 Yang, Z. et al. Recruitment of P-TEFb for stimulation of transcriptional elongation by the bromodomain protein Brd4. Mol. Cell 19, 535–545 (2005). 10.1016/j.molcel.2005.06.029

52 Flynn, R. A. et al. 7SK-BAF axis controls pervasive transcription at enhancers. Nat. Struct. Mol. Biol. 23, 231–238 (2016). 10.1038/nsmb.3176

53 Ohno, S. I. et al. Nuclear microRNAs release paused Pol II via the DDX21-CDK9 complex. Cell Rep. 39, 110673 (2022). 10.1016/j.celrep.2022.110673

54 Blagosklonny, M. V., Krueger, B. J., Varzavand, K., Cooper, J. J. & Price, D. H. The Mechanism of Release of P-TEFb and HEXIM1 from the 7SK snRNP by Viral and Cellular Activators Includes a Conformational Change in 7SK. PLoS ONE 5 (2010). 10.1371/journal.pone.0012335

55 Chao, S.-H. et al. Flavopiridol Inhibits P-TEFb and Blocks HIV-1 Replication. J. Biol. Chem. 275, 28345–28348 (2000). 10.1074/jbc.C000446200

56 Baumli, S. et al. The structure of P-TEFb (CDK9/cyclin T1), its complex with flavopiridol and regulation by phosphorylation. EMBO Journal 27, 1907–1918 (2008). 10.1038/emboj.2008.121

57 Czudnochowski, N., Bosken, C. A. & Geyer, M. Serine-7 but not serine-5 phosphorylation primes RNA polymerase II CTD for P-TEFb recognition. Nat. Commun. 3, 842 (2012). 10.1038/ncomms1846

58 Oksuz, O. et al. Transcription factors interact with RNA to regulate genes. Mol. Cell 83, 2449–2463 (2023). 10.1016/j.molcel.2023.06.012

59 Ozes, A. R., Feoktistova, K., Avanzino, B. C., Baldwin, E. P. & Fraser, C. S. Real-time fluorescence assays to monitor duplex unwinding and ATPase activities of helicases. Nat. Protoc. 9, 1645–1661 (2014). 10.1038/nprot.2014.112

